# Losing the split ends: unraveling the mechanisms of fiber branching during blood clot formation

**DOI:** 10.64898/2026.10.13.758846

**Authors:** Aravind Elangovan, Dylan M. Miller, Can Cai, Key Crosley, Heather A. Belcher, Rachel M. Lee, Chad M. Hobson, David M. Hart, Martin Guthold, Brittany E. Bannish, Nathan E. Hudson

## Abstract

The coagulation cascade generates a three-dimensional fibrin gel that seals wounds and prevents the loss of blood. Extensive research has defined the structural and mechanical characteristics of fibrin gels and the various biochemical, physiological, and pathological factors that alter those properties. However, there has been no direct visualization of fibrin polymerization with the spatial and temporal resolution to characterize the fibrin branching process. Thus, a mechanistic explanation for the origins of fibrin structures remains unresolved, with several hypothesized models for how fibrin branching occurs. Using SimView light-sheet microscopy, we captured videos of fibrin fibers forming junctions and gelling. Results: We observed four distinct pathways for fibrin fibers to interact and bind, all predicated on the observation that tumbling and diffusing fibers collide and stick together. Branching occurred due to the binding of mature fibers, not through the splitting of fibers or through the lateral association of small fibrin oligomers (protofibrils), as had been previously hypothesized. Diffusion simulations of rod-like fibers confirmed the branching mechanisms, the relative frequency of the various binding pathways, and the distribution of fiber angles after binding. Scanning Electron Microscopy (SEM) imaging also corroborated the relative frequency of various types of fibrin junctions. Moreover, SEM enabled the visualization of junctions at different timepoints, demonstrating that additional fibrin strands adhere to the initial binding site, strengthening the interaction. These results establish a mechanistic framework for fibrin branching and gel formation and define an innovative experimental approach for future interrogations of fibrin polymerization.

## Introduction

Fibrin gels form the structural and mechanical backbone of blood clots. Polymerization is initiated when thrombin cleaves fibrinopeptides A and B (FpA, FpB) from fibrinogen, revealing knobs ‘A’ and ‘B’, which bind with corresponding holes ‘a’ and ‘b’ in nearby fibrin molecules. This interaction results in the formation of half-staggered two-molecule thick oligomers, commonly referred to as protofibrils. Protofibrils then bundle together and laterally aggregate to form fibers, which often feature a distinct substructure consisting of multiple interwoven fibrin strands (likely consisting of protofibrils or bundles of protofibrils).(1, 2) Finally a 3-D branched gel is formed, which is further reinforced through the covalent crosslinking of fibrin by FXIIIa.(3, 4) Despite extensive study of fibrin polymerization (1, 2, 5–10), the mechanisms by which fibrin branching occurs remain unresolved, with at least seven distinct proposed models (Supplemental Figure 1).(6, 7, 11–14) Since the branching mechanisms govern the eventual 3-D architecture, resolving these processes is critical for understanding the origins of altered fibrin structures observed in many diseases.(15–17)

Light scattering studies during polymerization have resulted in at least three competing models of fibrin branching (junction formation) (Supplemental Figure 1). Some studies argued that junctions originate from a rooted tree where fibrin fibers undergo restricted branching as they grow due to the fibrin molecule acting as a copolymer. (8, 14, 18) Other studies proposed that as long protofibrils diffuse spatially and rotationally, they will eventually bind to other protofibrils and become immobilized in a disordered network.(7, 19, 20) A more recent hypothesis suggests the formation of a transient, ‘Y-ladder’ intermediate oligomer, followed by a transition to a protofibril structure, with incorrectly binding monomers generating branches.(6) Collectively these models differ in whether branching is an intrinsic feature of protofibril growth or arises from independent protofibril interactions.

Electron microscopy (EM) studies of fibrin junction structures led to additional hypotheses on how branching occurs. Complicating these results is the observation that fibrin polymerizes into a fine mesh of mostly thin protofibrils at high pH and salt concentrations and a coarse gel of thicker fibers at physiological pH and salt conditions, so polymerization mechanisms may differ under these two environments, and some networks consist of both thin and coarse fibers.(1, 21) In fine protofibril meshes, formed at high pH and ionic strength, both symmetric and asymmetric protofibril junctions were observed, suggesting two formation pathways: 1) “tetramolecular” junctions, with one branch being twice the width of the other two, formed by the lateral association or bifurcation of protofibrils,(1, 11, 22) or 2) trimolecular junctions, with all three branches of equal width, originating from the misincorporation of a monomer in a growing protofibril, resulting in a branching site.(11, 22) In contrast, coarse networks formed under physiological salt and ionic strength often exhibited aligned 23 nm molecular banding (corresponding to half the length of a fibrin molecule) across all branches of a junction, supporting a mechanism in which branching arises from fiber strand (protofibrils or bundles of protofibrils) bifurcation that maintains molecular lateral registry (Supplemental Figure 1).(2, 10) Protofibril twisting was hypothesized to limit the number of inter-fiber contacts thereby promoting the formation of branches. Observations of fibrin strand convergence into, and separation from, “cables” (thick fibers consisting of multiple strands) led to the additional hypothesis that “anastomosis” between growing fibers forms a branched 3D network.(1) Mechanical studies indicating that junctions are comparable in strength to fibers and also exhibit cross-struts preventing branch point failure align with this view.(23, 24) In sum, static snapshots of junctions from EM provide a detailed view of branching structures, leading to hypotheses that junctions form from either the convergence or bifurcation of fibrin protofibrils/strands or the misincorporation of molecules in a protofibril.

While most studies describe a predominance of three-branch junctions, (25–29) four (or more)-branch junctions increase in frequency under specific conditions including low ionic strength (50 mM NaCl).(27) Gel compression causes fibers to intersect and adhere leading to an increase in junctions with more than three branches .(25) Other reports of fiber “nodes”, where more than three fibers protrude from a single point, suggest alternative branching mechanisms, such as the twisting of fibers around each other or the nucleation of junctions by fibrin aggregates.(28, 29)

Attempts to visualize gelation using light microscopy have lacked the spatial or temporal resolution to observe branch-point formation.(30, 31) Early ultramicroscope studies reported short fast-moving rods jumping into and out of focus that consolidated into rigid needles.(32) Later investigations employing confocal or deconvolution microscopy identified mobile fibers at the early timepoints, followed by an increase in connectivity and junctions over time but were unable to resolve branching mechanisms.(31, 33, 34) Recent work has tracked fibrin formation induced by *Staphylococcus aureus*, however it is unclear if the gelation mechanisms are comparable to those induced by thrombin.(35)

Computational models have been developed to help understand the mechanisms and kinetics of fibrin branching. Early statistical models treated fibrin as tree-like polymers with fixed branch points.(14, 18) Other models implicitly represented branching through stochastic protofibril interactions or evolving fiber architecture.(36, 37) Several differential equations models treated branching as a consequence of the polymerization process, with individual monomers occasionally binding in non-linear configurations that initiate a new branch.(12, 13, 38) Coarse-grained molecular and particle-based simulations imposed bonding rules and hydrophobic interactions between fibrin monomers to reproduce branch formation (39, 40). However, most of these existing models either assume fixed topology or treat branching as a local chemical reaction, without explicitly modeling the dynamic interplay between fiber diffusion and physical contact.

Taken together, despite numerous modeling and experimental studies, it remains unclear whether the formation of fibrin junctions is governed primarily through mechanisms of protofibril growth or through interactions among diffusing fibrin polymers. Here, utilizing light-sheet microscopy (LSM), we observed polymerization with sufficient temporal and spatial resolution to visualize junction formation for the first time (Figure 1). We found that branching occurs primarily through collisions and adhesions between mature, growing fibers, rather than through protofibril splitting, and identified four distinct types of junction formation events. Scanning electron microscopy was congruent with these results and demonstrated a maturing fibrillar structure around junctions over time. Based on these results, we developed a new agent-based model incorporating fiber diffusion and contact-driven adhesion that reproduces key experimental features.

**Figure 1:**
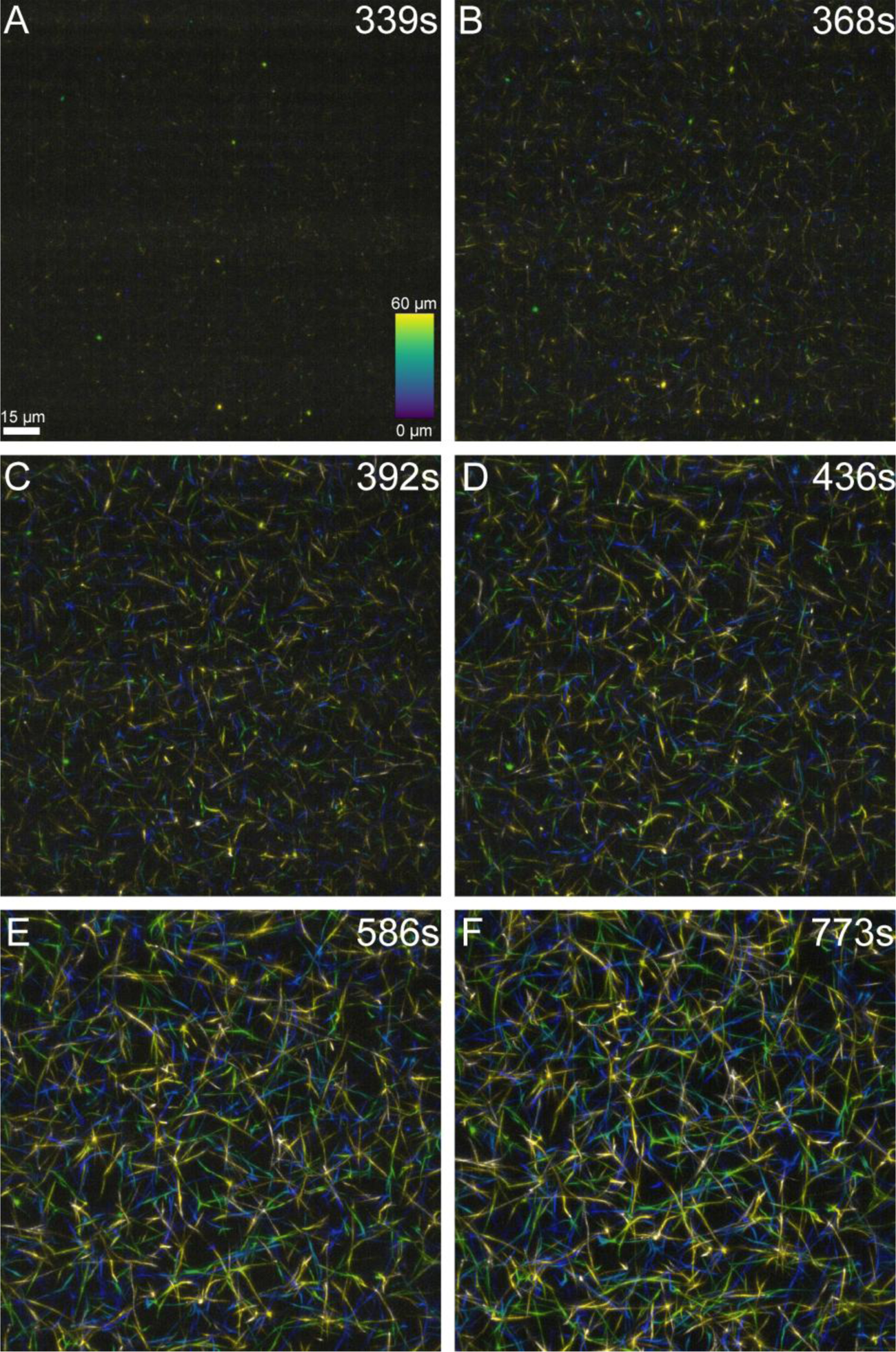
Time series of fibrin polymerization: A-B) Very short fibers appear, initially as diffraction-limited dots. C-D) Short fibers elongate and begin to bind. E-F) A stable and mature network develops. For all panels, the length scale is the same and the color scale represents fiber depth within the sample, with blue fibers being near the front of the volume and yellow near the rear. Z-depth colorized images were prepared using the Temporal-color Code feature in FIJI.(70)

## Methods

### Light-Sheet Microscopy (LSM)

A 60 µL human fibrinogen solution at 1 mg/mL was prepared by combining peak 1 fibrinogen (2.2 µL; 27.6 mg/mL; Enzyme Research Labs), AlexaFluor-488 labeled fibrinogen (8.9 µL; 0.15 mg/mL; Invitrogen, Carlsbad, CA), and HBS (HEPES-buffered saline; 48.9 µL; 20 mM HEPES, 150 mM NaCl, pH 7.4). A 60 µL human α-thrombin solution (Human α-Thrombin, Enzyme Research Labs) at 0.2 IU/mL was prepared by serial dilution of a stock thrombin vial at 339.3 IU/mL using HBS containing 10 mM CaCl_2_. Equal volumes (50 µL each) of fibrinogen and thrombin were mixed by gentle pipetting three times and briefly vortexed (<1 s), yielding final concentrations of 0.5 mg/mL fibrinogen and 0.1 IU/mL thrombin, with 2.2% AlexaFluor-488 labeled fibrinogen. The mixture was loaded into a 2.5 mm FEP (Fluorinated Ethylene Propylene; Zeus Inc., Orangeburg, SC) tube and immediately mounted for imaging.

Volumetric time series imaging (Figure 1; Supplemental Video 1) was performed on a modified Simultaneous Multiview (SiMView)(41) light-sheet microscope housed at HHMI Janelia Advanced Imaging Center. Volumetric images were acquired using a 40X 1.0 NA detection objective with 0.5 μm step size over 121 steps (60 μm depth) and 7 ms exposure time. After initial polymerization strands appeared, time series acquisition began at 2 s intervals and continued until intensity plateaued and polymerization had visually stopped (Figure 2).

**Figure 2:**
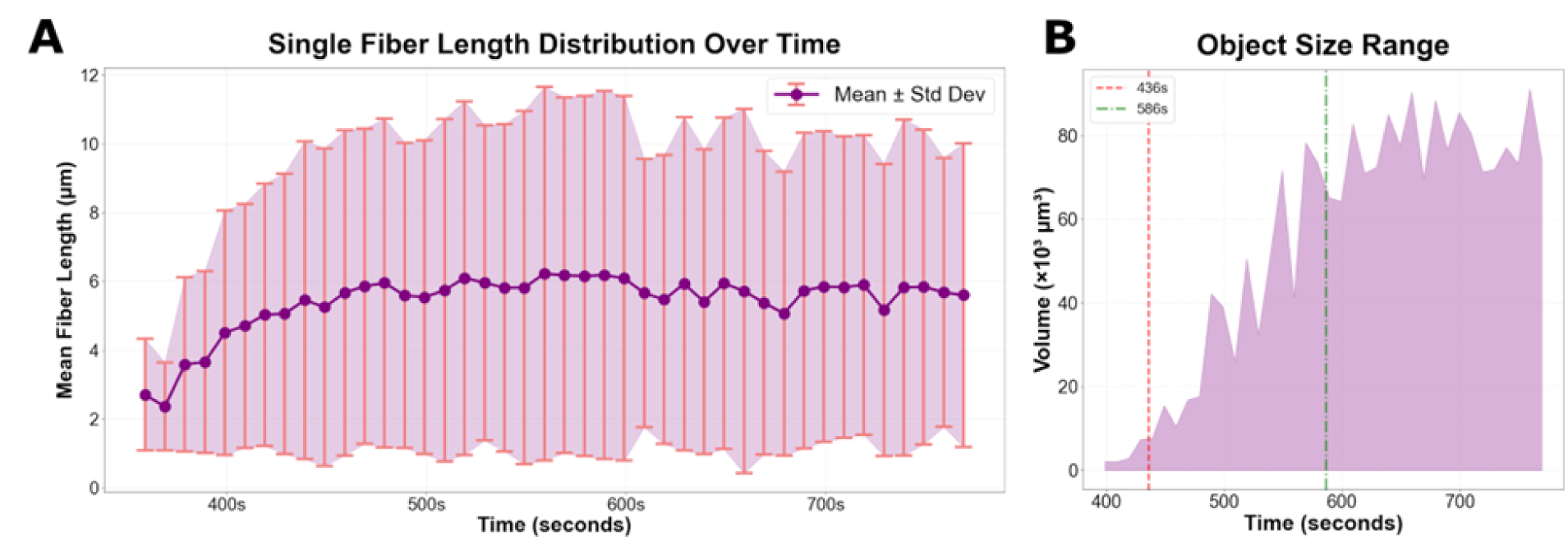
Length of unbound fibers, fluorescent object size, and number of objects over time in light-sheet microscopy. A) Quantification of the length of single, unbound fibers over time. B) Range of fluorescent object sizes at each timepoint. The largest volume datapoint represents the largest object, consisting of multiple connected fibers, at that timepoint, whereas the smallest volume object at the same timepoint consists of a single, small fiber. Annotated lines at timepoints 436 s and 586 s show the start of increase and leveling off respectively

### Imaris Analysis

All manual image analysis of LSM data was performed in three dimensions using Imaris 9.9 (Oxford Instruments, Abingdon, UK). Fibers were randomly selected by overlaying a 5×5 µm grid on the 165×165×60 µm³ volume at the final timepoint and selecting 30 fibers intersecting gridlines. These fibers were tracked across all timepoints, with those exiting the volume replaced. Junction formation type (orientation and binding mechanism of the interacting fibers), angles, and angle stability (consistency of angular orientation over time) were measured. Junction formation was identified at the timepoint where voxels of two fibers intersected (Figure 3; Supplemental Figures 2 and 3). Angles were measured using three points: one at the junction center and two along each fiber ∼1–2 µm away. Junction angles for T-, I-, and X-junctions were defined as the acute angle (maximum 90°), while L-junctions ranged from 0°–180° (Figure 4). Stability was assessed by measuring angles at 4 s intervals after formation, then at progressively increased intervals (∼8, 16, 64, and 128 s). Branch number per junction was determined by counting all branches within a 50×50×60 µm³ region.

**Figure 3:**
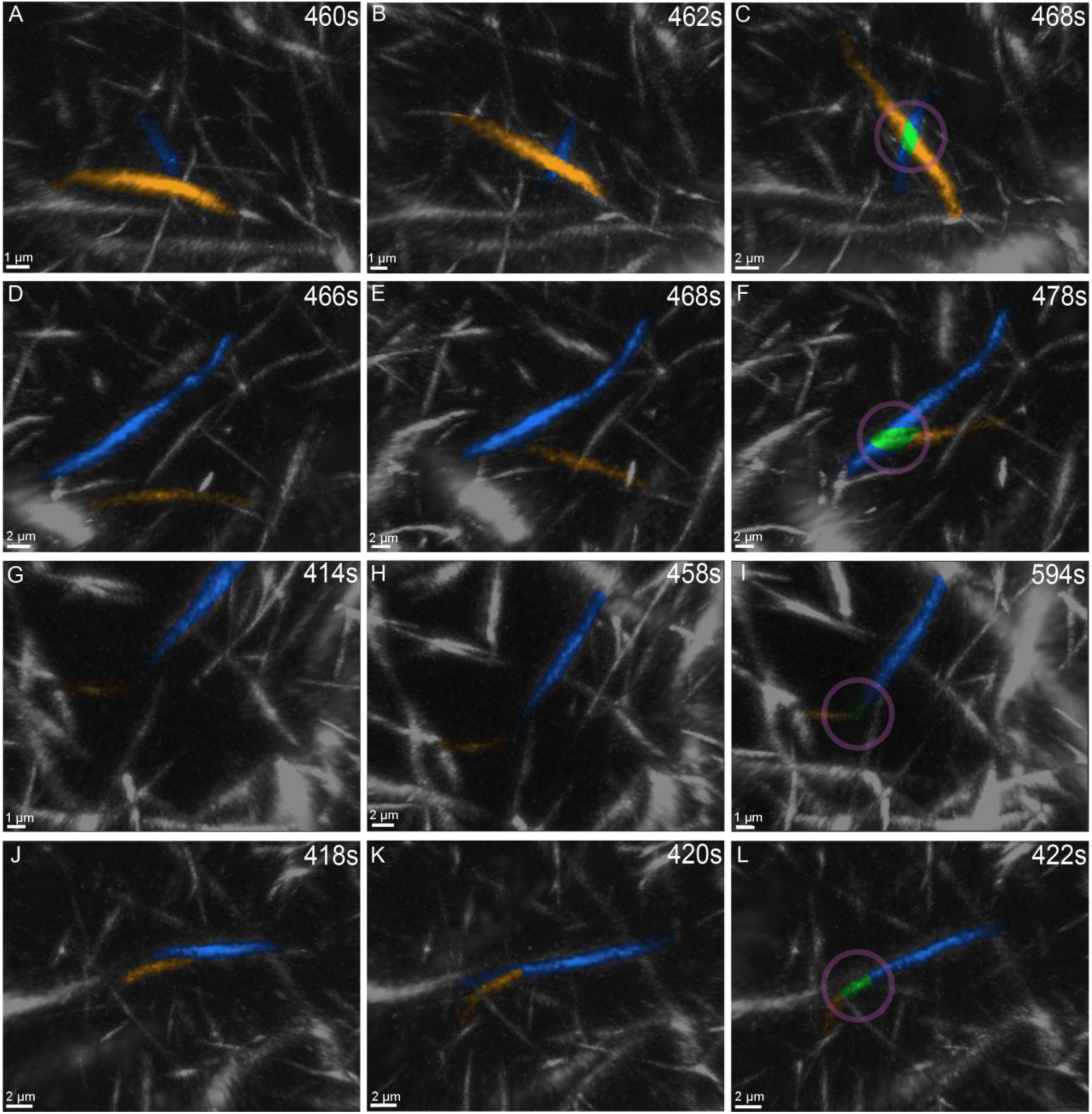
Time series of four types of junction formation events: In each row two distinct fibers that eventually bind are colored orange and blue. Fibers diffuse together in the left and center columns and bind to form a junction in the rightmost column. Junctions are highlighted by a magenta circle surrounding the fiber intersection point and green coloration of overlapping fiber segments. A-C) X-junction formation. X-junctions occur when the body of a fiber crisscrosses and binds to the body of another fiber. D-F) T-junction formation. T-junctions occur when the end of one fiber binds with the body of another fiber. G-I) L-junction formation. L-junctions occur when the end of one fiber binds with the end of another fiber. J-L) I-junction formation. I-junctions occur when the body of a fiber binds to the body of another fiber lengthwise and becomes one, longer fiber. Colorization to highlight the main structures was performed using colorized layers in Adobe Photoshop (Adobe Inc., San Jose, CA, USA).

**Figure 4:**
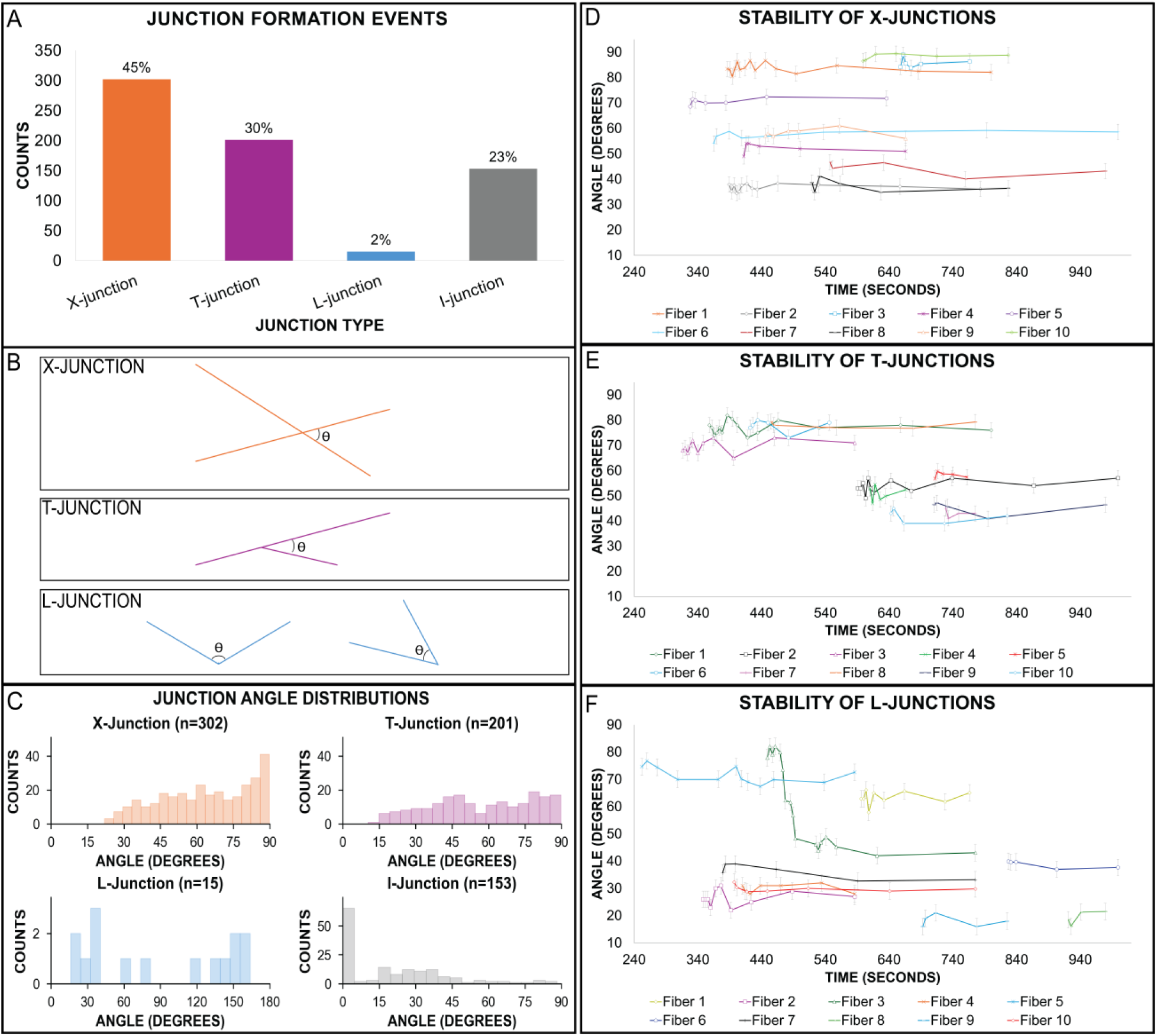
Distribution of junction formation event types and angles: A) Quantification of the number of junction formation events of each type observed in LSM (percentages shown above the bars). B) Definition of junction angles. X-, T-, and I-junction angles were all defined as the acute angle between the two binding fibers, while L-junctions could range from 0-180°, since an acute angle was not necessary for this type of junction. C) Distribution of binding angles for each junction formation event in LSM data, separated by type. Cumulative from n = 240 fibers from N = 8 biological replicates. D-F) Stability of junction angles. Junction angles for ten randomly chosen junctions of each type were tracked over time. The variance of nearly all junction angles was within ±5°, demonstrating angular stability after junction formation. Error bars reflect the estimated measurement error, derived from the standard deviation of 50 repeated measurements of the same angle.

### Quantitative Analysis

To complement manual analysis, automated quantification of 3-D fibrin networks was performed. Due to the failure of classical thresholding methods (Supplemental Table 1),(42) Indicator Guided Adaptive Thresholding (IGAT)(43) (Supplemental Figure 4) was utilized to segment thin fibers, followed by connected-component labeling (44, 45) and skeletonization.(46) Fiber length and object size were extracted from the skeletons and labeled objects using scikit-image.(47) To calculate junction branch number, skeletons were pruned(48) to remove spurious branches and merge erroneously split junctions.(43) However, the automated pipeline consistently undercounted higher-order junctions, particularly 4-branch nodes, and was therefore used as a supplementary measure (Supplemental Figures 5 and 6).

### Simulations

A 3-dimensional (3-D) agent-based model was developed in MATLAB (The Mathworks Inc., Natick, MA). Two-hundred-line segments of fixed length (6 μm, corresponding to average unbound fiber length at later timepoints; Figure 2), representing fibrin fibers, were randomly distributed in a 115 × 115 × 115 μm³ cube with reflecting boundary conditions. At each time step, fibers diffused rotationally and translationally according to corrected Broersma theory for rods(49), and fibers within 0.1 μm adhered to form junctions (Supplemental Figure 7). Joined fibers continued to diffuse at a reduced rate (diffusion coefficients scaled by 1/(number of attached fibers)³) with fixed connection angles. Simulations were run for 7 minutes of simulation time (Supplemental Figure 8). Model outputs included the number and type of junctions formed and the fiber angles at the junctions. Additional details are provided in the Supplement.

### Scanning Electron Microscopy (SEM)

SEM imaging of fibrin gels (peak 1 fibrinogen) followed the same preparation workflow as light-sheet experiments. Immediately after mixing fibrinogen and thrombin, 10 µL aliquots were transferred onto freshly cleaved mica. At defined time points (436, 586, and 773 s), reactions were quenched with 2 mL of 2% glutaraldehyde in sodium cacodylate buffer and fixed overnight. Samples were dehydrated by critical point drying (Tousimis SAMDRI-PVT-3D, Rockville, MD) and sputter-coated with gold (Cressington Model 108, Watford, UK).

A total of 232 SEM images were collected from nine samples (three biological replicates per time point) at magnifications ≥ 30,000X. Fiber diameter, junction length, and junction thickness (Figures 6 and 7; Supplemental Figure 9) were quantified in ImageJ using a randomized-grid sampling strategy adapted from a published protocol.(50) Junction thickness for 3- and 4-branch junctions was defined as the diameter of the thickest strand or cable (Figure 6 B&D).

**Figure 5:**
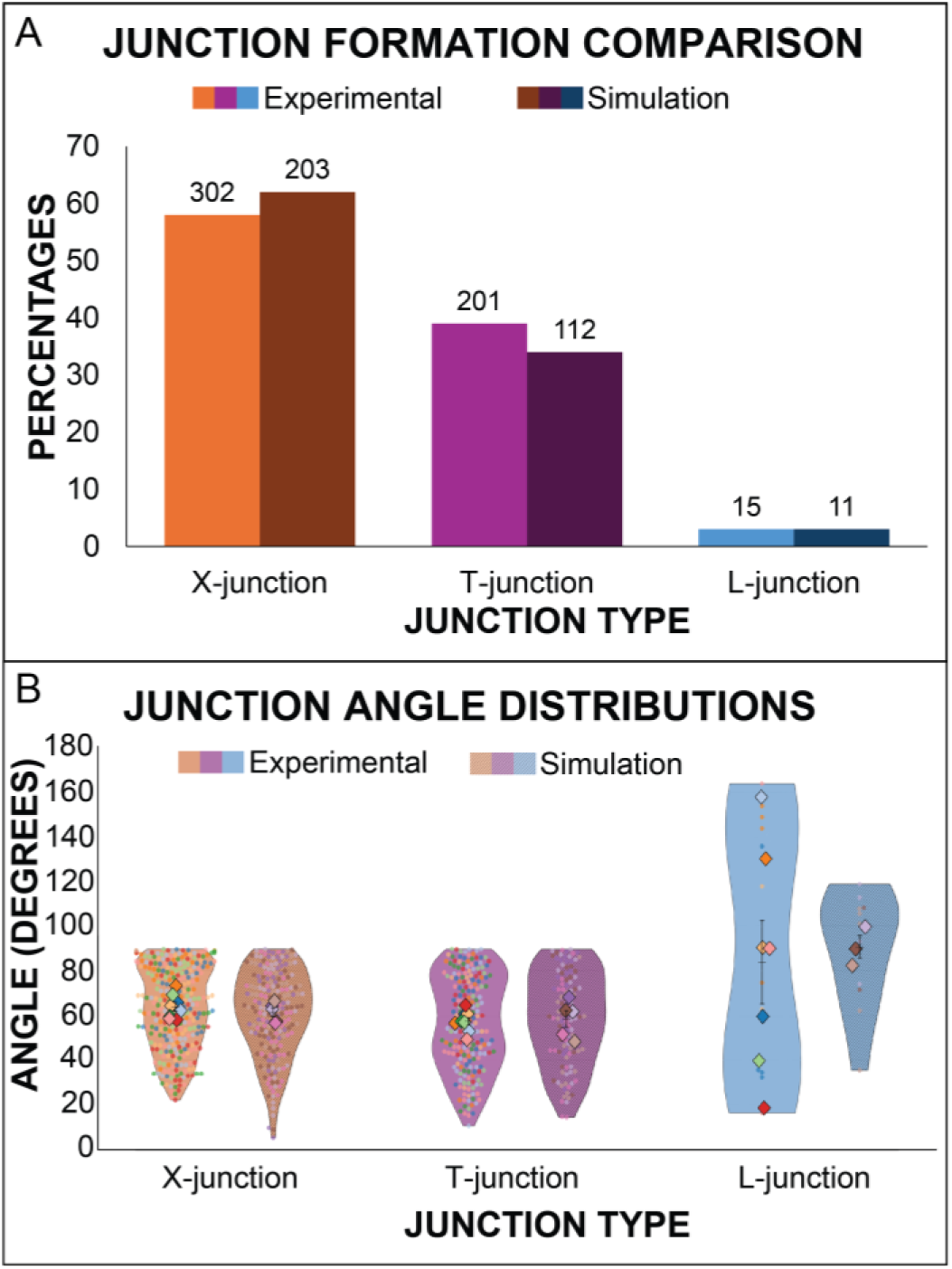
Distribution of junction angles in LSM and simulations: A) Comparison of the percentages of junction formation events of each type between LSM experiments and simulations. Because of their ability to rotate, I-junctions were excluded from this analysis. B) Comparison of angles from LSM (N=8 experiments) and simulations (N=5). P-values when comparing experimental and simulation angle distributions were greater than 0.05 for all junction types, when comparing both the combined data points or the mean values from each experiment/simulation suggesting strong congruence between the simulation and LSM data.

**Figure 6:**
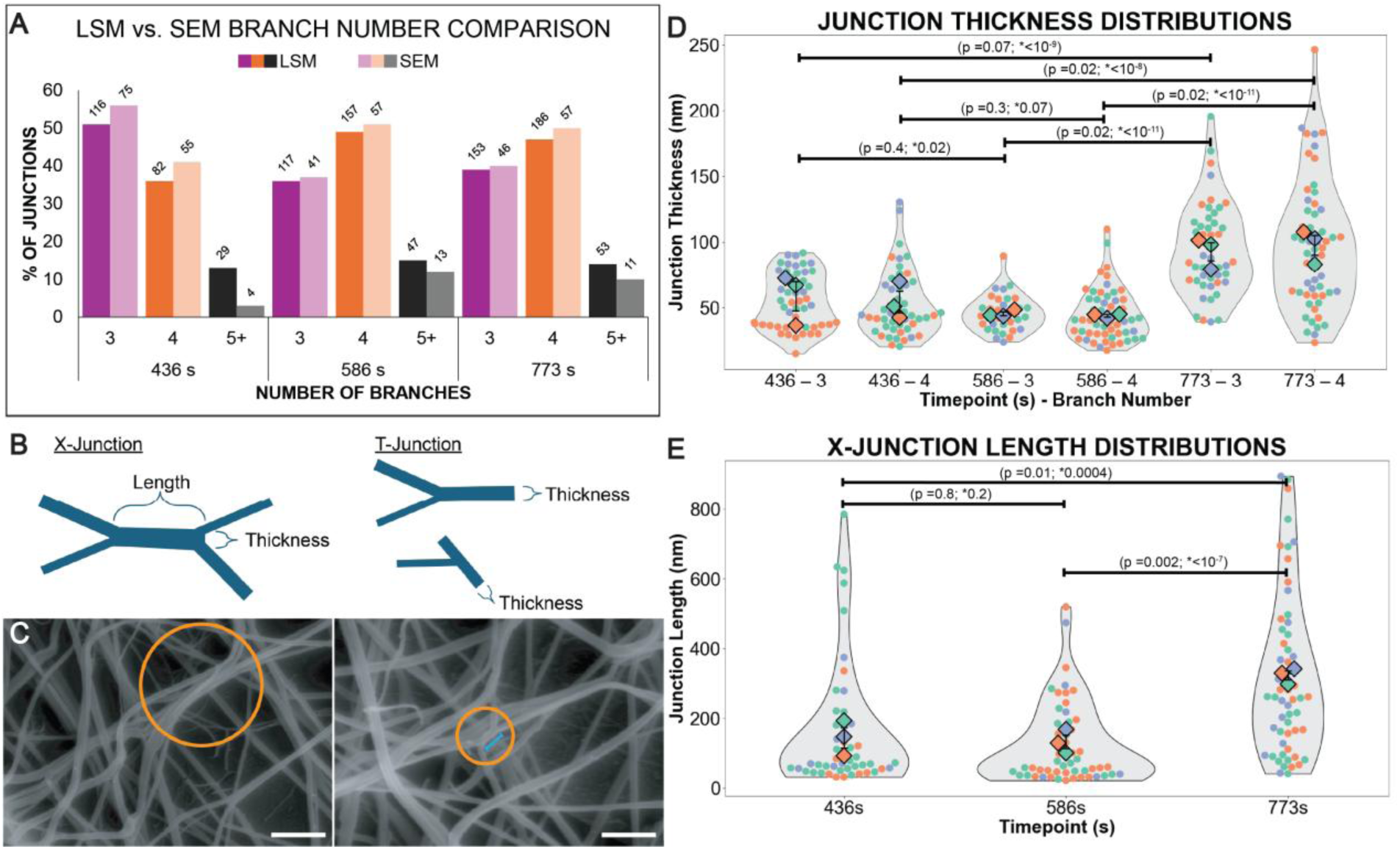
Maturation in branch type and branch structure over time: A) Quantification of the percentage of junctions with 3, 4, and 5+ branches at three different timepoints for both LSM and SEM data. N=3 experiments for LSM data and for each SEM timepoint, while n values are displayed above each bar. Multiple linear regression revealed no significant method-dependent temporal effect (Time × Method interaction: estimate ≍ 0, p = 1.00), indicating comparable temporal trends between LSM and SEM measurements. B) . Visual representation of X-junction and T-junction length and thickness. C) Scanning electron micrographs demonstrating fiber bundling and zippering at 773 s. Left: The orange circle highlights thick fiber bundles that converge and diverge. Right: Orange circle highlights a potential X-junction that has “zippered” together to form what look like two “T’s”. The junction length is demonstrated by the blue line. Scale bar: 1 µm. D) Superplots at each time point show the distribution of thicknesses for three-branch (T-junctions) and four-branch (X-junctions) junctions. Data for each timepoint represent N=3 biological replicates, with n=50 (436–3), n=33 (586–3), n=46 (773–3), n=49 (436–4), n=55 (586–4), and n=57 (773–4). E) Violin superplots of X-junction length distributions at three timepoints. Data for each timepoint represent N=3 biological replicates, with n=48 (436 s), n=54 (586 s), n=57 (773 s). The median length at 436 s (67.4 nm) and 586 s (59.9 nm) are comparable while the median length at 773 s (260.75 nm) is five times longer. Colored datapoints within each superplot indicate data from a specific biological replicate, with the mean value for each replicate being depicted as a diamond. P-values given for comparison between the mean values (left) and the combined data (*right).

**Figure 7.**
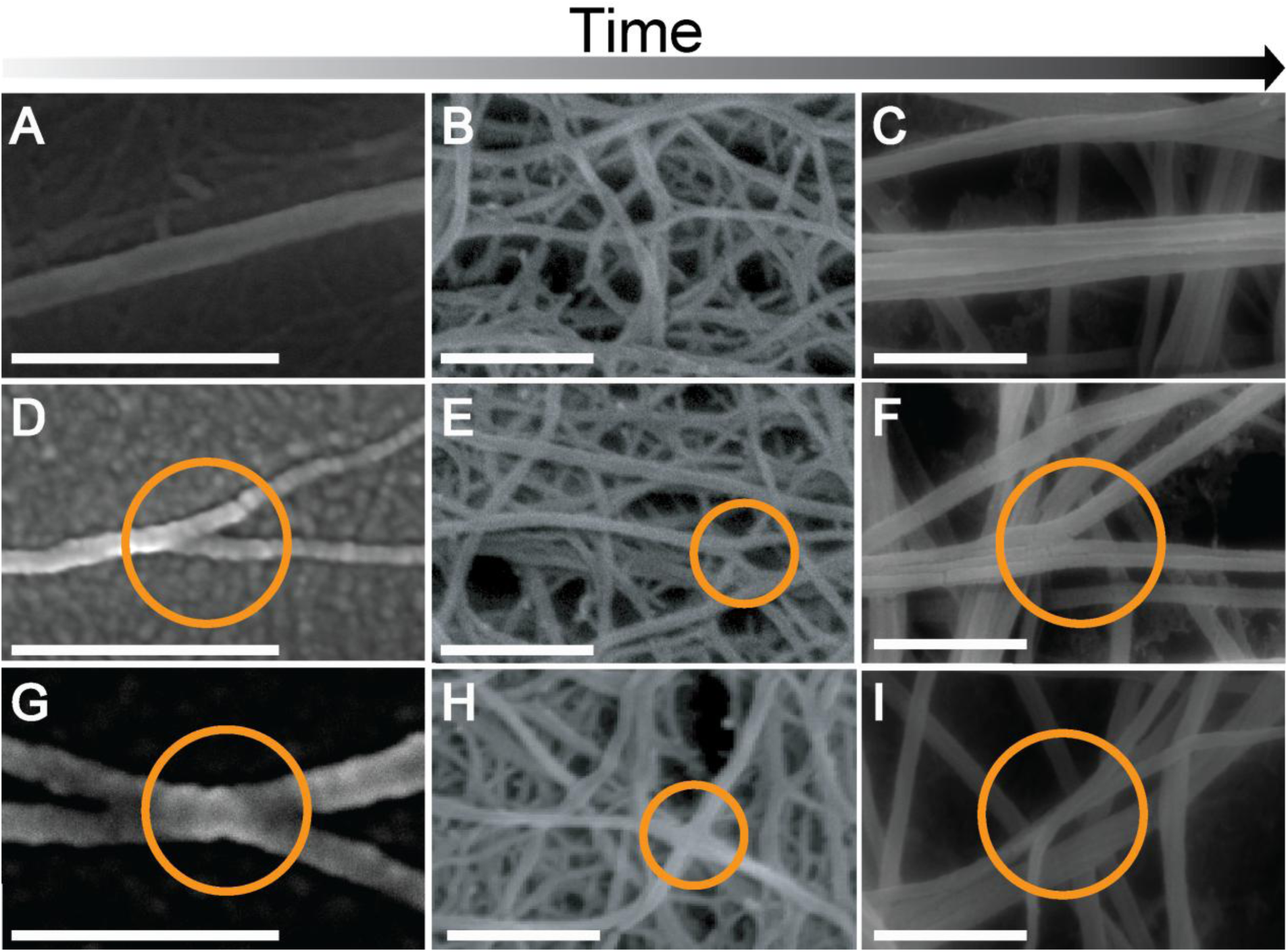
Representative SEM images of junction size and substructure during fibrin network maturation. Vertical columns correspond to timepoints of 436 s (left), 586 s (middle) and 773 s (right), respectively, while rows correspond to fibers (top), T-junctions (middle), and X-junctions (bottom). A) individual fibrin strand; B) incipient fiber fibers (cables), consisting of two or more distinct strands; C) mature fibrin fibers (cables), consisting of multiple strands; D) T-junction with single-stranded fibers; E) multi-stranded T-junction; F) T-junction with multi-stranded cables; G) X-junction with single stranded fibers. H) multi-stranded X-junction. I) X-junction with multi-stranded fibers (cables). Scale bar: 500 nm.

## Results

### Growth of Fibers

Automated identification of fibers in the LSM data demonstrated that fibers begin as small, diffusing objects that rapidly increase in length and size before binding to other fibers (Figure 2). The mean length of unbound fibers increased from 2 μm to 6 μm before leveling off ∼ 500s. The volume of the largest fluorescent object increased over time before plateauing ∼ 580 s, showing that fibers and clusters of fibers continued binding together until forming a relatively stable network structure.

### Tracking of Individual Fibers During Junction Formation

Thirty fibers from each of eight biological replicates (240 total) were manually tracked frame-by-frame during gelation. For each fiber, all junction formation events, including type and angle, were recorded. Four junction types were identified: “X”, “T”, “L”, and “I”, based on their resemblance to English letters (Figure 3). For example, T-junctions form when the end of one fiber binds to the middle of another, while X-junctions form from mid-fiber interactions, and L-junctions from end-to-end binding. I-junctions formed through multiple pathways (Supplemental Figure 3): 1) two fibers diffusing together to form a transient, angled bond prior to “snapping” into a single, straight fiber, typically within 10 s after the initial interaction (∼54%), and 2) two fibers diffusing together in parallel and overlapping partially or completely lengthwise before binding to form a single, straight fiber (∼40%). The straightness of I-junctions contrasts with L-junctions, where fibers remain at <180° after binding. Representative time series for all types of junction formation are shown in Figure 3 and Supplemental Videos 2–5. Notably, all junctions formed via diffusion-driven fiber interactions; no branching occurred via fiber splitting. Quantification of the number of each type of junction formation event (Figure 4A) shows X-junctions comprised 45% of events, followed by T-(30%) and I-junctions (23%), while L-junctions accounted for 2%.

In ∼6.5% (45/687) of binding events, an apparent junction was briefly formed, prior to the fibers separating, although it is unclear whether these reflect true binding or fiber proximity below the LSM resolution limit (∼320 nm). Of these fibers, 21 (3.1%) re-formed junctions, with a median unbound time of 8 s. Incomplete I-junctions, where the two parallel fibers did not completely adhere along the entire lengths, comprised 6% (9/153) of I-junction formation events (Supplemental Figure 3E-F,J-L), producing splayed fiber ends that resemble a T-junction, despite forming in a different manner. A rare transient event (observed once) involved two strands on one side of an X-junction zippering into a single fiber, creating a 3-branch T-structure.

### Junction Angle Distributions and their Stability

We quantified the initial angles of junction formation in the LSM data. X- and T-junction angle distributions peaked near 90°, with broad tails extending to ∼10° (Figure 4B-C). The I-junction distribution exhibited a sharp peak between 0°–7°, with a tail extending to 90°. Unlike other junctions, which maintain their initial angles (Figure 4), I-junctions can rotate into a parallel configuration, forming straight fibers due to the snapping mechanism described above. Simulations (Figure 5) agreed well with the X-, T-, and L-junction angle distributions but did not produce I-junctions, suggesting their formation requires mechanisms beyond random diffusion and collision, such as fiber bending, zippering along the fiber length, or post-binding snapping into place.

To assess junction stability in LSM, junction angles for 10 representative fibers were tracked over time (Figure 4D-F). Nearly all X-, T-, and L-junction angles remained stable for >100 s, deviating by only 5 ° - 10°. One L-junction changed its angle when a 3^rd^ fiber bound. I-junctions demonstrate a larger angular variance, as ∼54% formed initially at an acute angle, similar to a T- or X-junction, prior to snapping into a linear fiber (Supplemental Figure 3).

### Simulations

To verify junction formation mechanisms, the agent-based model (Supplemental Video 6) was run five independent times and results of these simulations were combined to create an *in silico* data set that contained 326 junctions. Simulation junction type percentages (Figure 5A) and angles (Figure 5B) were similar to experimental results for X-, T-, and L-junctions, indicating that diffusion and collision alone are sufficient to explain the formation of these junctions. I-junction formation, on the other hand, may require mechanisms beyond random collision, as discussed below.

## 3 vs 4-Branch Junctions Over Time

Since previous work has suggested a predominance of junctions with three branches,(25–27) we also quantified the number of 3-branch, 4-branch, and 5+-branch junctions in the LSM data at three time points (436 s [early in the junction formation process, represented by an increase in object size], 586 s [the time of nascent network formation, represented by a plateau in object size], and 773 s [mature gel has formed]) (Figure 6). Initially we attempted to do this using an automated approach similar to what had been used previously.(25, 43) Using this approach we found results of 89% 3-branch, 8% 4-branch, and 3% more-than-4-branches at the 773 s timepoint (Supplemental Figures 4 and 5). However, we compared these results to a subset of 393 fibers at the last timepoint of three samples that were quantified manually and surprisingly found that the automated approach significantly overestimated the number of 3-branch junctions (Supplemental Figure 5). Because of the discrepancy, we present the manually counted junction counts at three different timepoints (Figure 6A). Interestingly, the data show that 3-branch junctions are more common early, while 4-branch junctions are the most common species at the later timepoints.

### SEM Images of Junctions

To understand junction sub-structure, SEM imaging was performed on samples prepared under identical conditions to the light-sheet experiments. A total of 232 images were collected from nine samples (three biological replicates per time point) at 436, 586, and 773 s after polymerization initiation (Figure 6 and 7). Branch number comparisons (Figure 6A) showed good agreement between SEM and LSM data. At 436 s, we predominantly observed isolated fibers involved in single junctions (Figure 7A–C). Most fibers appeared as single strands; due to non-uniform thickness, diameters were measured at multiple positions, yielding a median of 34 nm (Supplemental Figure 9). The diameter of these strands suggests they consist of multiple protofibrils, though SEM resolution limited further detail. At later timepoints, mature fibers formed cable-like structures composed of multiple strands within 3-D networks. By 773 s, median fiber diameter increased to 62 nm, while individual strands within cables measured 27 nm.

We also interrogated the structures of the junctions themselves using SEM. At 436 s, both T- and X-junctions could be observed, typically consisting of intersecting single fiber strands (Figure 7D,G). While the 436 s branches consisted of single fiber strands, at the 586 and 773 s timepoints, branches often consisted of multi-stranded fibers (cables) (Figure 7E-F,H-I). To quantify these differences, we measured the thickness of branchpoints at different time points (Figure 6 B-D). Early timepoint branches (436 s and 586 s) had median thicknesses ∼ 40 nm for both X- and T-junctions, whereas at 773 s, the thicknesses were ∼95 nm, suggesting that junctions mature and grow wider over time, particularly after fibers begin to connect into a polymerized network. Moreover, at 773 s, X-junctions often demonstrated a “stretched X” shape (Figure 6 B, C&E,), where the X-junction looked like two T-junctions separated by a length. Because of this, we assigned all “double T” junctions as X’s if the length between bifurcations was less than 1000 nm (∼3X the resolution limit of LSM) for comparing SEM and light-sheet junction counts (Figure 6A). At early timepoints (436 s and 586 s), the median X-junction length was ∼60 nm, while at 773 s, the median length was ∼260 nm (Figure 6E).

## Discussion

The results presented here provide the first real-time visualization of fibrin junction formation. Tracking 240 fibers as they polymerize using LSM clearly demonstrates that fiber junctions form almost exclusively through the binding of diffusing and tumbling fibers, rather than the splitting of protofibrils within growing fibers. The mean length (∼ 5 μm; Figure 2) and diameters (36 ±12 nm; Supplemental Figure 9) of the fibers at the time of early junction formation (436s), indicates that they are mature fibers rather than oligomers or protofibrils,(34) and rules out most hypotheses related to branching occurring during protofibril growth under physiological salt concentrations (Supplemental Figure 1).(6, 12, 19, 51)

X-junction formation events were the predominant species observed in tracking the branching of individual fibers, with simulation results corroborating their critical role in the gelation process. Adhesion of diffusing fibers that collide together is consistent with previous observations that fibrin is “sticky” and prone to bind to other fibers and materials.(52, 53) Intriguingly, the number of 4-branch junctions increased over time (Figure 6A), suggesting that X-junctions may be a function of fiber length, such that longer fibers are more likely to form X-junctions, while shorter fibers are more likely to form T-junctions. Because local fibrinogen and thrombin concentrations control fiber growth rates and final fiber lengths,(54, 55) we hypothesize that clotting conditions favoring quickly forming longer fibers will exhibit more X-junctions.

T- and L-junctions form via similar mechanisms, depending on where the ends of the two colliding fibers adhere. In the LSM data, the formation of both T- and L-junctions appeared to terminate fiber growth at the binding site (e.g., in the T-junction in Figure 3 D-F, when the end of the “orange” fiber binds to the middle of the “blue” fiber, the “orange” fiber stops growing longitudinally at that end; in the case of an L-junction, the adhered ends of both fibers stop growing). On the other hand, formation of an X-junction did not terminate the growth of either fiber, enabling fibers to continue growing in length after a binding event (Supplemental Figure 2). Thus, T- and L-junctions play an important role in preventing fibers from continually growing and spanning the entire length of the network (Supplemental Figure 2), which accounts for the absence of free fiber ends in mature fibrin gels.(27, 56)

SEM images of junctions at different timepoints provide important details on junction maturation and, when combined with the light-sheet data, force a reinterpretation of previous electron microscopy analysis of junction structures. T-junctions at early time points (436 s) showed single stranded fibers forming junctions, while at later timepoints the junctions were thicker and consisted of multi-stranded cables (Figure 6D and Figure 7). Previous literature has commonly interpreted multi-stranded junctions, and the twisting of fibrin strands at mature junctions(1), as occurring through the splitting of fibers or as the coalescing of parallel protofibrils.(2, 11, 51, 57) However, our data suggest that multi-strandedness of fiber junctions likely occurs through an initial binding event (X-, T-, etc.) followed by the later addition of new strands, since we did not observe any junctions forming through the splitting of fibers in the LSM data. This reinforcement mechanism likely explains the appearance of cross-struts when junctions are stretched.(23) In addition, the observation of stretched X-junctions in SEM data, particularly at the 773 s timepoint (Figure 6C,E and Figure 7I), suggests that strands may zipper together over short distances after X-junction formation (Figure 8A). Because these junction lengths are commonly near the ∼320 nm LSM diffraction limit (Figure 6C), zippering would be difficult to detect by fluorescence imaging but readily observed by electron microscopy, potentially leading to the overcounting of 3-branch junctions (Supplemental Figure 4). Future experiments, perhaps using super-resolution microscopy,(15, 56) to track junction development are warranted, to validate these results in hydrated fibers and avoid potential artifacts introduced through the SEM drying process..

**Figure 8:**
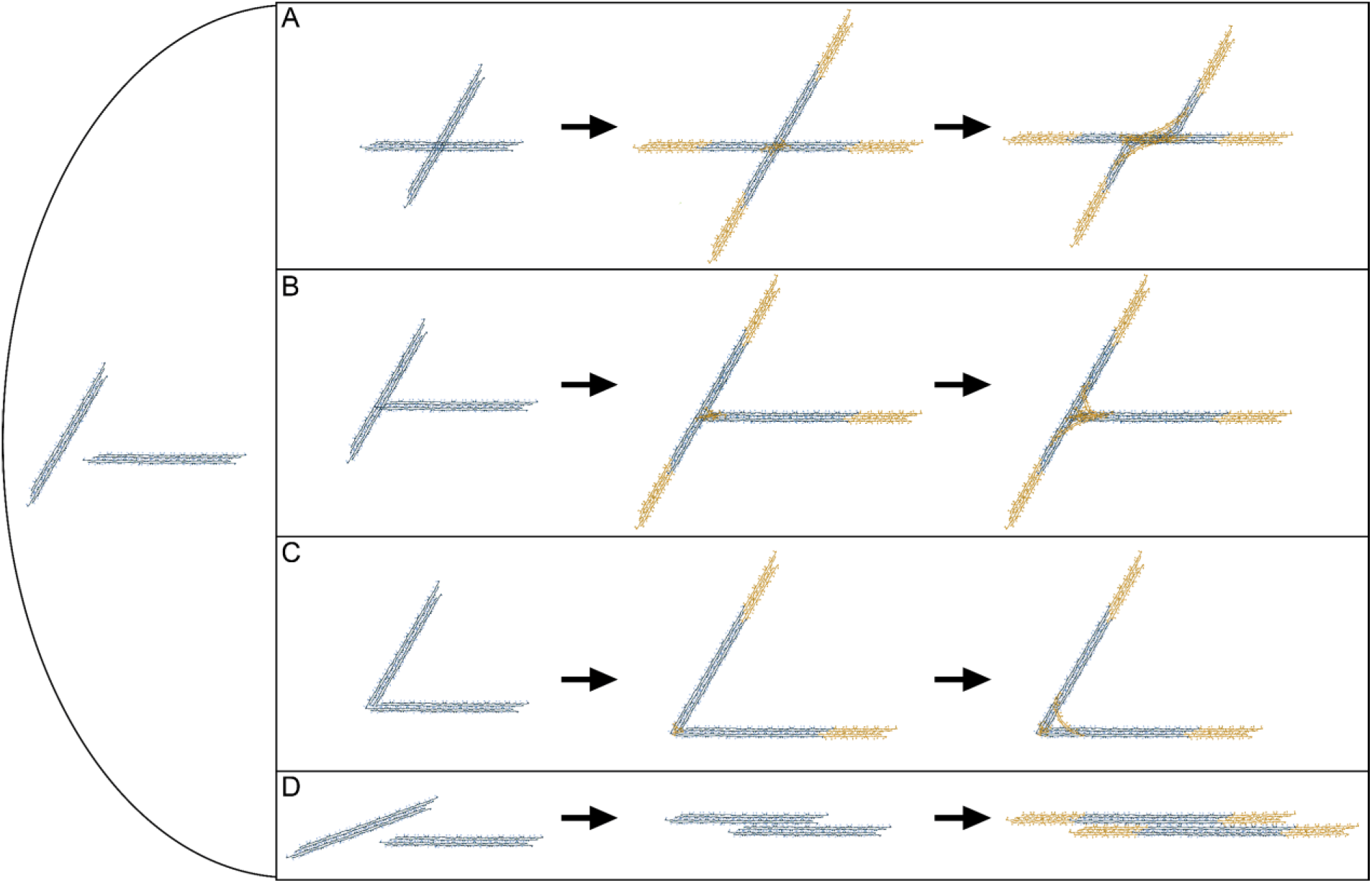
Model of junction formation and maturation process. Cartoon depicting A) X-B) T-, C) L-, and D) I-junction formation from two fibrin strands. Yellow objects depict protofibrils that bind to an already formed junction to strengthen the adhesion or grow the fiber length. Also depicted in (A) is an X-junction that zippers together over a short length.

Junction angle distributions from the LSM data are relatively flat and match simulations, indicating no preferred formation angles (Figures 4C and 5B) and suggesting that the mechanisms responsible for the initial binding interaction are non-specific. Additionally, the angle of X-, T-, and L-junctions remains consistent over time in the LSM data (Figure 4D-F), suggesting that the initial binding interactions, despite being non-specific, are strong enough to maintain the bond during diffusion, even prior to any reinforcement from later binding strands. In the low Reynolds number environment, viscous drag dominates inertia, which may help to maintain the proximity of the fibers, even when adhesion is weak.

I-junctions are defined primarily by their end-product, which is a single, longer and thicker straight fiber. We hypothesize that when I-junctions bind and then snap into place the initial bond is not strong enough to fix the angle, so the two fibers rotate into a parallel configuration where the numerous adhesion sites between protofibrils (2, 51) fasten the final I-junction configuration (Figure 8 and Supplemental Figure 3). It is intriguing that while X-, T-, and L-junction angles remain stable after the initial binding event, I-junctions can rotate. These distinctions likely arise from differences in the molecular architecture of the binding sites, and future high-resolution studies of fiber adhesions could resolve these features. The lack of I-junctions in simulations likely arises from the fixing of angles upon fiber collision and concurs with LSM data that the diffuse-and-bind mechanism alone cannot account for I-junction formation. Future simulations should include a fiber “snapping” mechanism. Interaction potentials that promote fiber alignment may also be important, as few simulated junction angles occur at < 10°.

A variety of biochemical, cellular, and biophysical factors are known to affect the final gel structure, including fibrinogen and thrombin concentrations,(26, 55, 56, 58) calcium and buffer composition,(26, 59) FXIIIa crosslinking,(60) plasma composition,(61–63) cellular content,(64) shear rates,(65, 66) and pathological conditions,(15, 17) thus there are myriad effects that could alter branching. For example, “fine clots” (formed at pH ≥ 8.5 or salt concentrations ≥ 450 mM) exhibit both trimolecular and tetramolecular junctions, (11, 22) suggesting that branching mechanisms under those conditions may be different than in the “coarse” clots of this study. Future studies should understand the influences of these factors on branching mechanisms. As a proof-of-concept, polymerization data was collected using LSM at a 6X higher fibrinogen concentration (3 mg/mL) to interrogate whether a higher density of fibrinogen would alter branching. Interestingly, the junction formation processes were the same, but the distribution of X-, T-, L-, and I-junctions differed (Supplemental Figure 10), with a statistically significant shift away from I-junction formation and towards X-junctions. We hypothesize that the diffuse-and-bind mechanism of junction formation will account for fibrin branching under most conditions, due to the ubiquitous nature of fibers growing and diffusing regardless of the local biochemical and cellular environment.

LSM and SEM provide a complimentary experimental platform for interrogating junction formation and structure, nonetheless limitations should be considered. LSM cannot resolve whether fibers are truly adhering or merely closer than ∼320 nm in distance. These spatial limitations are mitigated by the temporal resolution, which allows the ability to follow fluorescent objects over time. The diffusion constants of fibers (*D_parallel_* ≈ 0.3725 μm^2^/s, see supporting information), suggest that if fibers are not bound together, they will diffuse apart (beyond 320 nm) within a few seconds (1-3 frames). Thus, when fibers remain together for longer than this period, we can confidently conclude that they have bound together.

On the other hand, SEM has high spatial resolution but cannot follow the time evolution of a single fiber. SEM is subject to some network collapse during dehydration and drying, which may make fibers and junctions appear thinner. Nonetheless, Electron Microscopy has provided the basis for much of the previous structural characterization of fibrin fiber junctions,(1, 2, 50) making it a useful technique for comparison in this study. Moreover, recent protocols (followed in this report) have been developed that limit network collapse during SEM dehydration(50), and studies comparing fiber diameters between superresolution microscopy techniques and SEM suggest limited fiber shrinkage in SEM, particularly at the lower thrombin concentrations used in these studies.(56, 67) Additionally, even if SEM results in slightly reduced fiber dimensions, the sample preparation steps are unlikely to systemically change two key observations: 1) the observed number of strands in junctions (junction thickness, Figure 6D and 7F&I) or 2) the number of fiber branches meeting at a junction—that is, to convert a T-junction into an X-junction or vice versa (Figure 6A). Thus, the main SEM-related observations of this paper should be independent of artifacts from sample preparation. One significant advantage introduced by SEM in these studies is the ability to resolve the substructure of fibrin junctions at different timepoints (Figures 6 and 7). Although the bifurcation of fibrin strands at junctions has been noted previously,(1, 11, 27) the observation that the junction thickness (average number of strands) increases over time (Figure 6D) is an important clue in understanding junction maturation. Uncovering the mechanisms of fibrin branching has significant pathophysiological implications as fibrin architecture directly determines the mechanical stability, permeability, and susceptibility to breakdown of a clot.(4, 57, 68) Highly branched, dense networks exhibit reduced permeability and increased resistance to fibrinolysis, contributing to thrombotic disorders such as ischemic stroke,(69) whereas loosely branched clots are prone to lysis and bleeding.(17) Leveraging an understanding of how fibrin branching occurs thus has the potential to provide mechanistic insights into the origins of aberrant pathological clot structures and guide the development of therapeutic strategies that target fibrin formation and stability.

In conclusion, fibrin arrives in its 3-D branched structure primarily through the diffusion, collision, and adhesion of mature, growing fibers. After initial adhesion events, junctions are reinforced by the binding of further fiber strands at the site. These results fundamentally reorient our understanding of how fibrin networks form and provide a framework for future studies interrogating fibrin polymerization mechanisms under pathological conditions.

## Acknowledgements

We thank Taylor Dement and Marcus Case for their contributions to this project. This work was supported by the Howard Hughes Medical Institute (HHMI) through an Advanced Imaging Center proposal. We thank all the members of the Advanced Imaging Center at HHMI Janelia Research Campus for their support while we were using the SiMView microscope and the subsequent data analysis. Research reported in this publication was supported by the National Heart, Lung, And Blood Institute of the National Institutes of Health under Award Number R15HL148842 (NEH, subaward BEB and MG) and by the National Science Foundation under Award Numbers 2332977 (BEB) and 2332976 (NEH). The content is solely the responsibility of the authors and does not necessarily represent the official views of the National Institutes of Health nor of the National Science Foundation. BEB received support from the University of Central Oklahoma College of Mathematics and Science CURE-STEM award. DMM was supported by an East Carolina University Undergraduate Research and Creative Activity (URCA) award and an Undergraduate Research in Physics (URPA) award. The funders had no role in the study design, data collection and analysis, decision to publish, or preparation of the manuscript.

## Authorship Contributions

A.E., D.M.M., H.A.B, D.M.H., C.M.H, and R.M.L. collected and analyzed the light-sheet microscopy data, C.C. collected and analyzed the scanning electron microscopy data, K.C. wrote the simulation code and analyzed the data, M.G., B.E.B., and N.E.H. supervised the project. N.E.H., A.E., and D.M.M. wrote the manuscript. All authors discussed the results and implications and commented on the manuscript prior to submission.

