## Supplemental Data for "Losing the split ends: unraveling the mechanisms of fiber branching during blood clot formation"

### Expanded Methods:

All LSM and SEM experiments were performed at final fibrinogen and thrombin concentrations of 0.5 mg/mL and 0.1 U/mL respectively except for the LSM data in Supplemental Figure 10, which used a final fibrinogen concentration of 3 mg/mL. Preparation of these solutions is described in detail below. In all experiments a stopwatch was started at the moment of initial mixing and used to quantify the start of data collection on the instrument.

#### Light-Sheet Microscopy (LSM) Experimental Details

AlexaFluor-488 fibrinogen preparation

The initial concentration of AlexaFluor-488-labeled fibrinogen (Invitrogen) was at 1.5 mg/mL, which was diluted to 0.15 mg/mL by mixing 2 µL of fibrinogen solution with 18 µL of HBS (HEPES Buffered Saline; 20 mM HEPES, 150 mM NaCl, pH 7.4). After dilution, the 20 µL AlexaFluor-488-labeled fibrinogen solution was centrifuged at 13,000g for 15 minutes to remove aggregates prior to adding it to the final fibrinogen solution.

Fibrinogen solution preparation

Peak 1 human fibrinogen was used to prepare the fibrinogen solution. To prepare the 60 µL fibrinogen solution, 2.2 µL of peak 1 fibrinogen (Enzyme Research Labs; stock conditions of 20 mM Tris, 150 mM NaCl, pH 7.4) at 27.6 mg/mL was mixed with 8.9 µL of the AlexaFluor-488 fibrinogen (0.15 mg/mL) solution, and 48.9 µL HBS (HEPES Buffered Saline). The final concentration of fibrinogen, for all experiments except the ones clearly labeled as 3 mg/mL, before mixing, is 1 mg/mL, with 2.2% AlexaFluor-488-labeled fibrinogen molecules. For experiments with a final fibrinogen concentration of 3 mg/mL, a similar approach was followed, but the final concentration before mixing was 6 mg/mL, while still retaining 2.2% AlexaFluor-488 fibrinogen, with volumes needed to reach that concentration being adjusted accordingly.

Thrombin solution preparation

The initial concentration of 2 µL of human α- thrombin (Enzyme Research Labs; stock conditions of 3107 NIH u/mg and 3.58 mg/mL in 50 mM sodium citrate/200 mM NaCl/.1% PEG-8000/pH 6.5)) was at 339.3 IU/mL. Thrombin activity was calibrated in IU/mL using a colorimetric assay (Thrombin Activity Assay Kit ab234620; Abcam, Cambridge, UK). Then 18 µL of HBS+CaCl2 (HBS with 10 mM Calcium Chloride) was added to dilute the concentration to 33.9 IU/mL. 2 µL of thrombin at 33.9 IU/mL was transferred to a new centrifuge tube and diluted to 3.4 IU/mL by adding 18 µL of HBS+ 10 mM $CaCl_2$. Next, 3.5 µL of thrombin at 3.4 IU/mL was transferred to a new centrifuge tube with 56.5 µL of HBS+CaCl2 bringing the final concentration of thrombin, before mixing, to 0.2 IU/mL.

Microscopy

For light-sheet excitation, a 488 nm pulsed laser (PicoQuant VisUV-488; 40 MHz pulse frequency; 50 µW entering the excitation objective) was expanded into a static light-sheet via a Powell lens (Laserline Optics; LOCP-8.9R05-3.0); the non-expanded axis was matched in beam size to the original design of the SiMView microscope.(1) A custom designed water-dipping 6.4x illumination objective (Special Optics; 54-12.5-31) was used to focus the light-sheet onto the sample. Emission was collected by a 40x 1.0 NA water dipping objective lens (Zeiss; 421462-9900-0000), filtered through a bandpass emission filter (Semrock; FF01-540/80), and focused onto a detector (Hamamatsu Orca Flash 4.0 v3). The fields of view (FOV) used was 1024 x 1024 pixels at 0.1625 µm per pixel for a total size of 166.4 x 166.4 µm. The Multiview imaging capability was not used in this context; that is, single-sided excitation and single-sided detection were used. Volumetric images were acquired using a 40X 1.0 NA detection objective with 0.5 µm step size over 121 steps (60 µm depth) and 7 ms exposure time. After initial polymerization strands appeared, time series acquisition began at 2 s intervals (including the time to acquire the image stack) and continued until intensity plateaued and polymerization had visually stopped. Following acquisition, single z-stacks were collected at five unexposed FOVs as controls for light-induced polymerization defects.

#### Statistical Methods

Data "Superplots" were created in accordance with recommended guidelines.(2) P-values used to assess difference between the data distributions in the Superplots throughout the text were calculated using Welch's t-test in MS Excel and were calculated for both the mean values of the biological replicates as well as the combined data. Unless otherwise noted, a significance threshold of $\alpha = 0.05$ was applied to all statistical tests.

Fisher's exact tests were performed in R(3) using two-sided tests with the built-in fisher.test() function; Bonferroni correction was applied to pairwise comparisons, yielding an adjusted significance threshold of $\alpha = 0.0125$. For each combination of method (LSM or SEM) and time point, the fraction of observations in each branch number category was calculated by dividing the raw count by the total count across all branch number categories. A multiple linear regression was then fit to these fractions as a function of time, method, and branch number, including two-way interactions between time and both method and branch number (lm: Frac ~ Time × (Method + Number)), using R's built-in lm() function.

### Branching Mechanisms

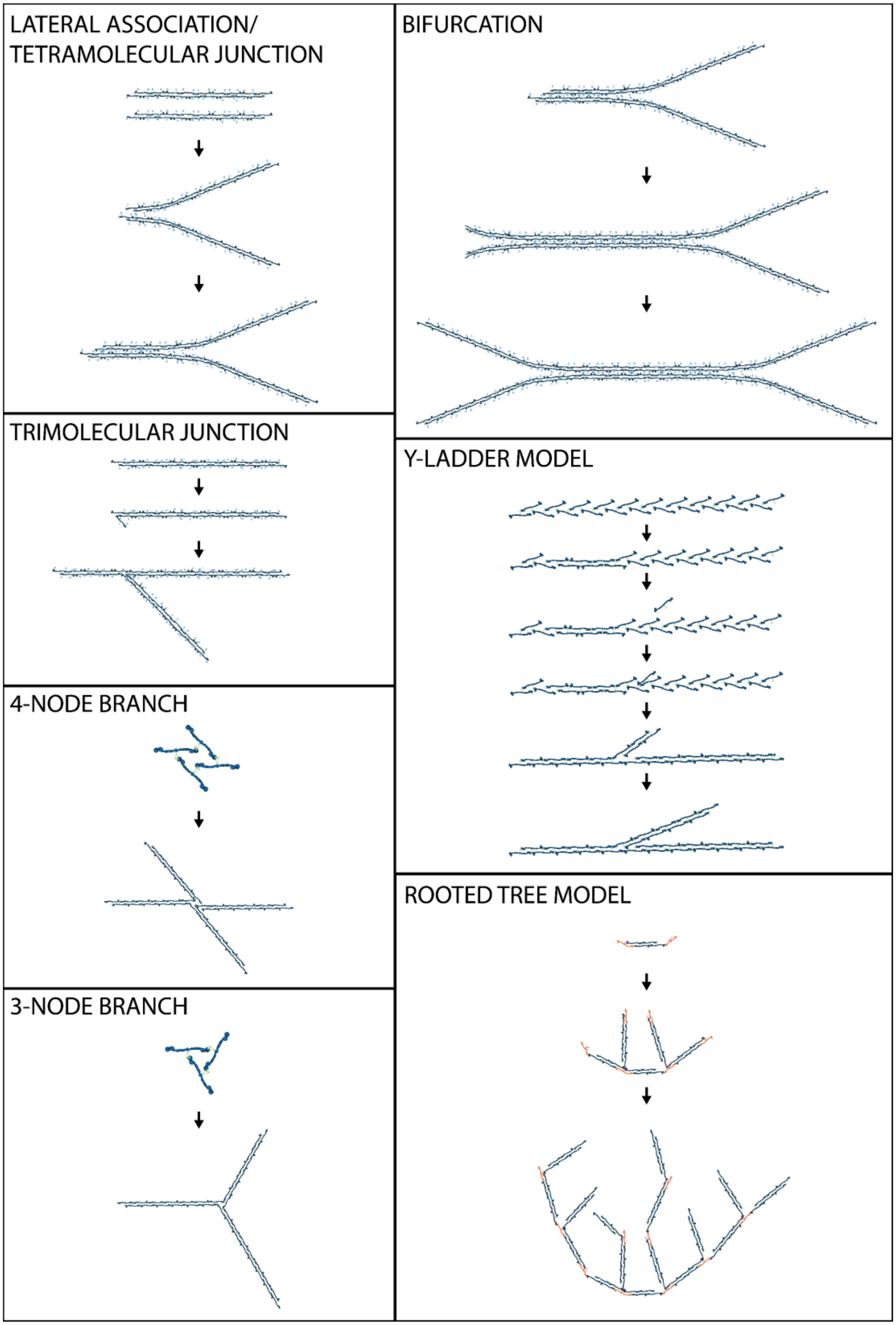


**Supplemental Figure 1: Current hypothesized models of junction formation.**

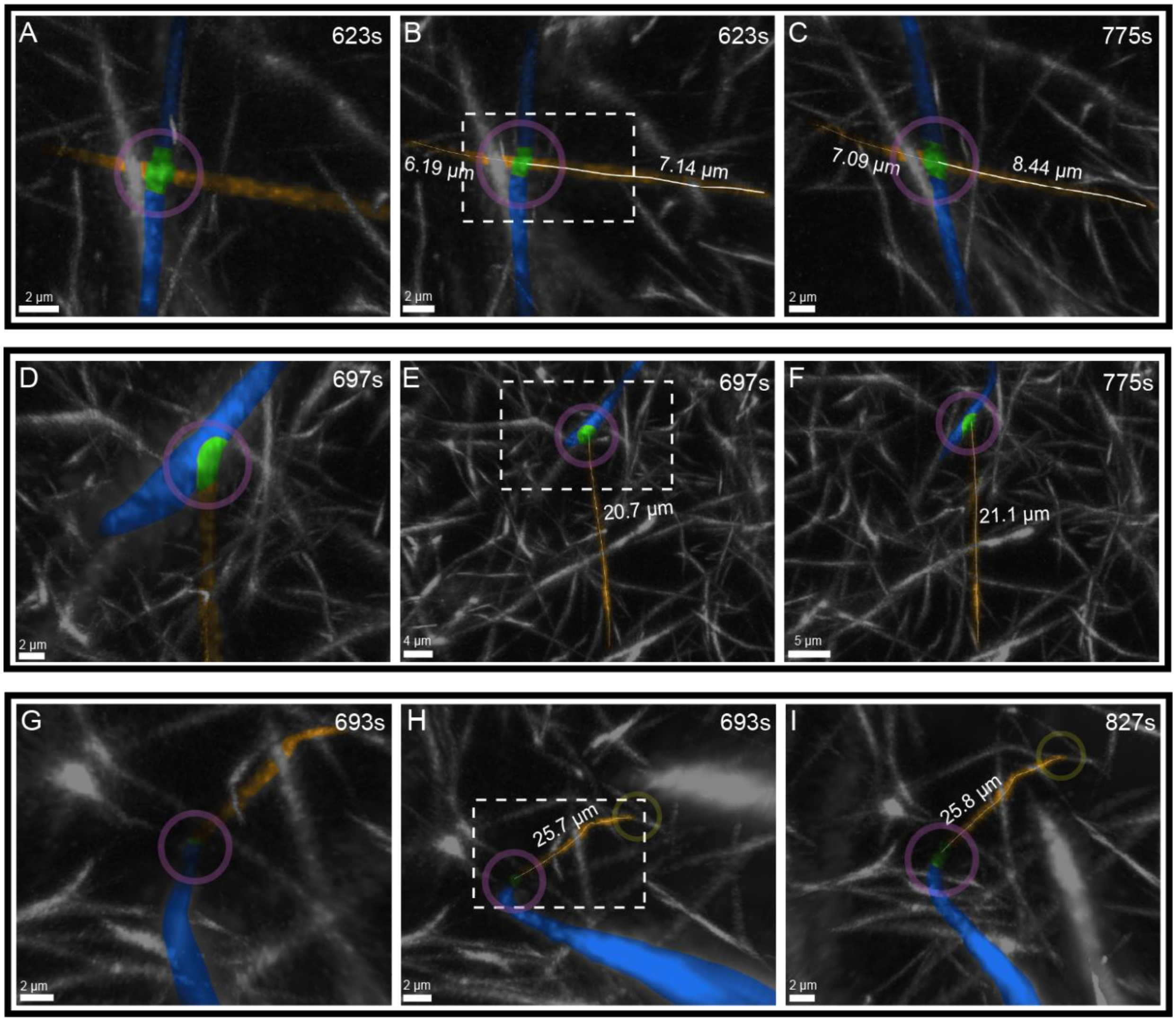


**Supplemental Figure 2: Growth/Terminality of fibers after junction formation.** In each row two distinct bound fibers are colored orange and blue with the intersection point highlighted with a magenta circle and green coloration of overlapping fiber segments. A-C) X-junction formation and fiber growth. A) Zoomed in X-junction image corresponding to the dashed rectangle in B), of an X-junction that just formed. B) Zoomed out view of the same junction as A) but with the entire fiber length shown. Representative growing fiber traced with a white line. C) X-junction at later timepoint. Both ends of the fiber continue growing after junction formation. D-F) T-junction formation and fiber growth. D) Zoomed in T-junction image corresponding to the dashed rectangle in E). E) Zoomed out view of the same junction as D) but with the entire fiber length shown. Representative fiber traced with a white line. E) T-junction at later timepoint. The unbound end of the fiber continues growing after junction formation, while the fiber end at the junction stops growing longitudinally. G-I) L-junction formation. G) Zoomed in image corresponding to the dashed rectangle in H), of a L-junction that just formed. H) Zoomed out view of the same junction as G) but with the entire fiber length shown. Representative fiber traced with a white line. I) L-junction at later timepoint. One end of the fiber forms an L-junction, the other end of the fiber forms a T-junction, thus this fiber has a fixed length and no longer grows longitudinally.

#### I-junction Formation

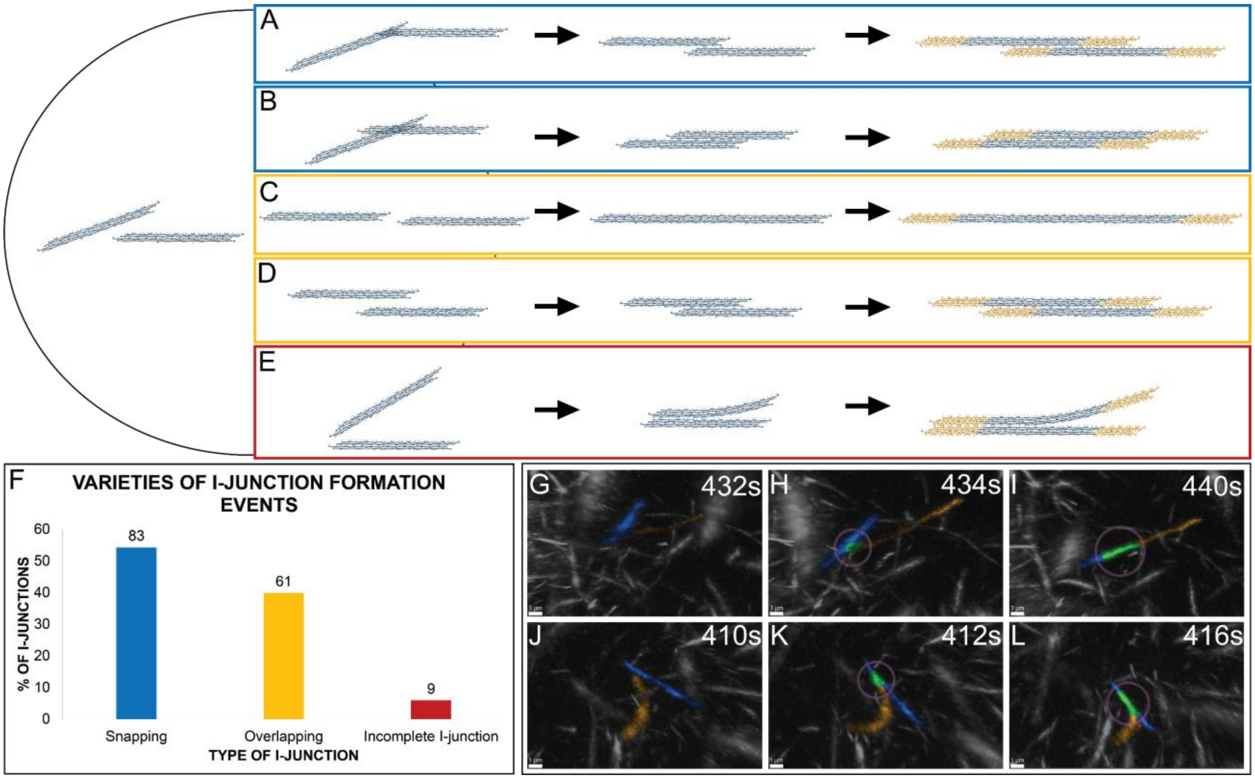


**Supplemental Figure 3: Varieties of I-junction formation events**. A-E) Cartoons of I-junction formation events. Yellow objects depict protofibrils that grow the fiber length after the initial binding event. A-B) I-junction formation through fiber snapping into place. Fibers stick at an angle and then snap into a straight configuration prior to binding. C-D) I-junction formation through binding of aligned fibers. Fibers align in parallel and then overlapping strands bind together. E) An incomplete I-junction that partially forms, but the strands don't fully bind, leaving a T-shaped junction. F) Percentages of I-junctions that form via the three varieties observed (raw counts above bars). G-I) Experimental time series of an I-junction forming through binding and snapping into place. J-L) Experimental time series of an incomplete I-junction. Two distinct fibers that eventually bind are colored orange and blue. Fibers diffuse together in the left and center columns and bind to form a junction in the rightmost column. Green coloration represents overlapping fiber segments.

### Quantitative Analysis

Quantitative analysis begins with the binarization of raw light-sheet data into analyzable segments. Several classical methods of binarization were tested for their usefulness with these data, but all failed to both handle the thin fibrin fibers and resolve constituent branches of junctions accurately. Methods included both local thresholding techniques like adaptive gaussian and global methods like OTSU.(4)

Instead of these methods, an approach more tailored to the thin structures that fibrin forms was needed. This was found in the form of Indicator Guided Adaptive Thresholding (IGAT).(5) This approach first applies anisotropic diffusion followed by a multi-scale vessel enhancement in the form of a Frangi Filter.(6) P-points are selected from the Frangi response; a density threshold of p-points then determines ‘indicator’ points of high p-point density. Finally, for each indicator point, a local adaptive thresholding is computed in the window surrounding that point.

This binarization was compared to hand-annotated ground truth data and a variety of common binarization methods. Prospective methods were tested by their similarity to the ground truth set by Jaccard Similarity, F1 score, and precision. Tested methods include an adaptive gaussian, simple Frangi filter, global OTSU, and IGAT. The results are summarized in Supplemental Table 1.

**Supplemental Table 1: Binarization comparison results**

| Method | Jaccard Similarity | F1 Score | Precision |
|---|---|---|---|
| IGAT | 0.3203 | 0.4852 | 0.3385 |
| Adaptive Gaussian | 0.2226 | 0.3641 | 0.2279 |
| Global OTSU | 0.1699 | 0.2904 | 0.1715 |
| Frangi Filter | 0.1383 | 0.2431 | 0.1424 |

After binarization, individual objects were labeled based on 3-dimensional connected components implemented in the connected-components-3D Python library.(7, 8) These connected objects were then pruned based on size, with objects less than 125 voxels in volume removed, to suppress noise while retaining small fibers.(9) While this provides the connected regions within the volume, fibers connected in a junction are labeled as a single object. Thus, to analyze the fibers that make up a given object, objects underwent skeletonization according to the Lee method(10) as implemented in the sci-kit image Python library (Supplemental Figure 4).

To further analyze these skeletons, methods from the IGAT analysis were employed.(5) These methods attempt primarily to address the pruning of false junctions and junction merging. Three main sources of errors are spurious junctions along the length of fibers, segmentation erroneously connecting physically disconnected objects, and the widening of 4-branch junctions into multiple 3-branch junctions. Skeletons are cleaned through pruning, starting with a simple step of removing junctions whose branch-to-endpoint length is under that of the average fiber thickness. Following this initial step aimed at smaller perturbations, the intensity profile of branches is analyzed. Quadratic regression analysis is run on the intensity across the fiber, and branches whose profile follows a bright → dim → bright (or ‘peak → valley → peak’ as Chen et al describe), are removed as false branches.

Despite these methods, there was a discrepancy in the branch number of junctions analyzed (Supplemental Figure 5). Likely due to the width of 4-branch junctions, many 4-branch junctions are split into separate junction points that are each 3-branch (Supplemental Figure 6). The solution attempted as part of previous analyses is as follows: candidate pairs are identified by closeness of two branch points. Candidate pairs within the thickness of thin fibers are automatically merged. Candidates outside of this distance are considered for merging based on their branch orientations. For each candidate pair, each branch is approximated by a line using regression fitting. The centroid of each branch segment is calculated, and the distance from one branch centroid to the other branch line is computed. Finally, the angle between the two lines is calculated. If both distances are less than the fiber thickness, and the angle is under 20 degrees, the two branches are considered colinear and merged.

Quantitative analysis of branch numbers calculated in this fashion conflicted with hand counts. It falsely identified 3-branch junctions as the majority of junctions after gelation when hand counts showed 4-branch junctions were the true majority at that timepoint. This was reflected in both light-sheet hand counts and SEM imaging. This discrepancy may have been due to several reasons, which include the splitting of 4-branch junctions into two, 3-branch junctions, binarization near junction points causing over or underestimations of connectivity to other fibers (and pruning steps not entirely accounting for this), or binarization resulting in split fibers.

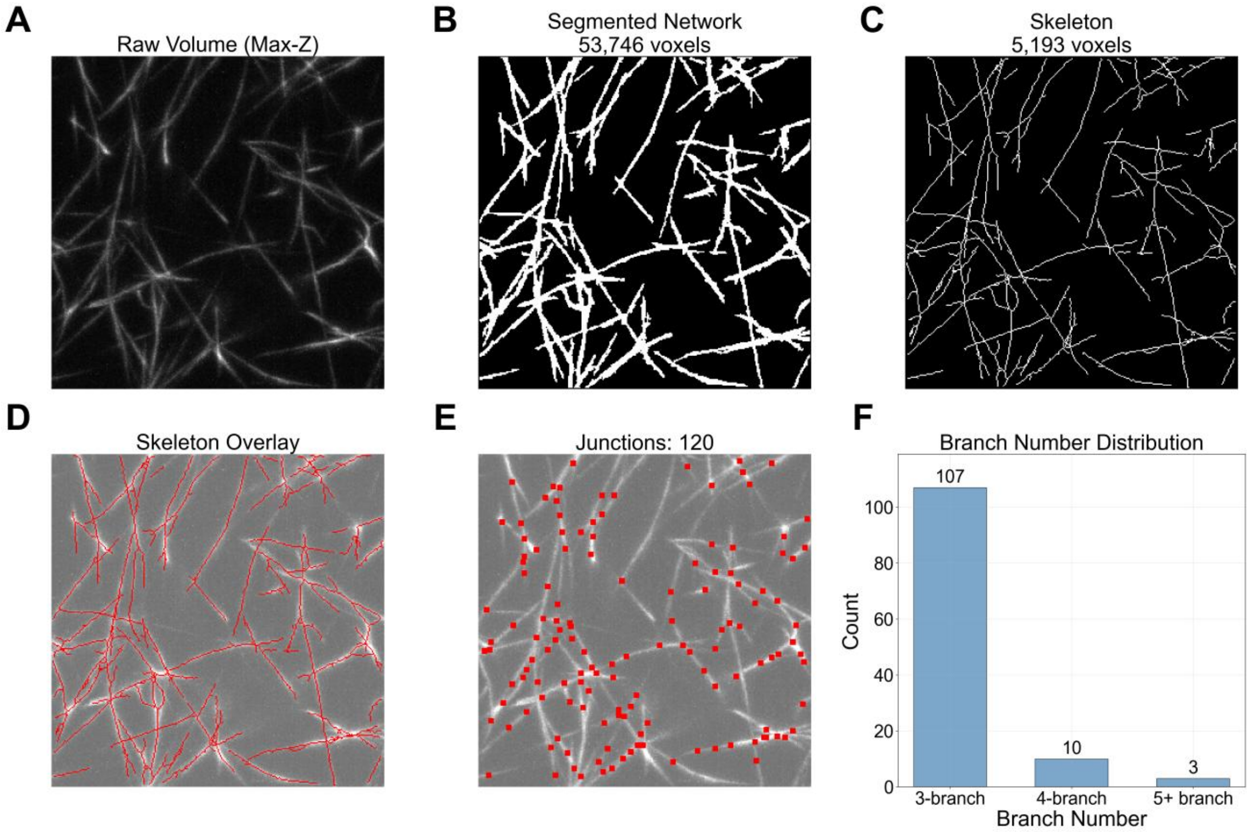


**Supplemental Figure 4: IGAT segmentation and skeleton analysis.** A) Max z-projection of central region of a volume with raw intensity. B) Segmented max z-projection of same central region via IGAT. C) Skeletonized max z-projection. D) Skeleton overlayed onto raw data. E) Junction points identified from the skeleton overlayed onto raw data. F) Branch number distribution measured from identified junction points. Each panel is the same 48.3 x 48.3 x 24.8 μm sub-volume.

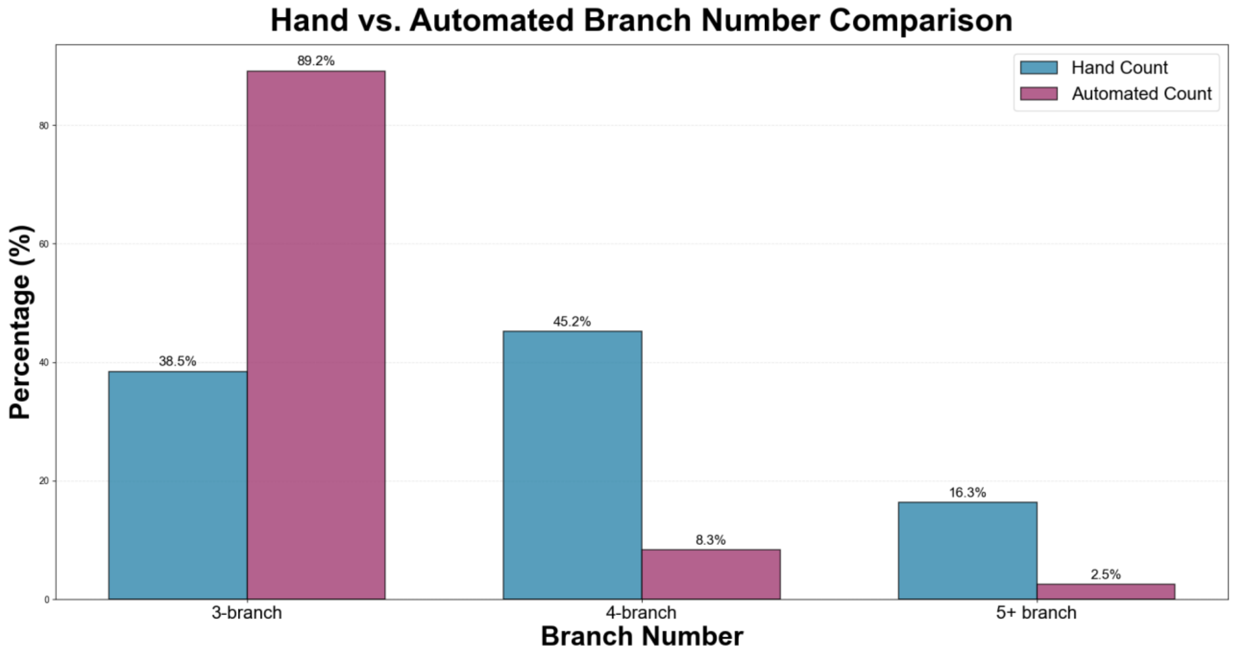


**Supplemental Figure 5: Comparison of automated and manual branch number counts.** Quantification of junctions with 3, 4, and 5+ branches using either hand or automated counts.

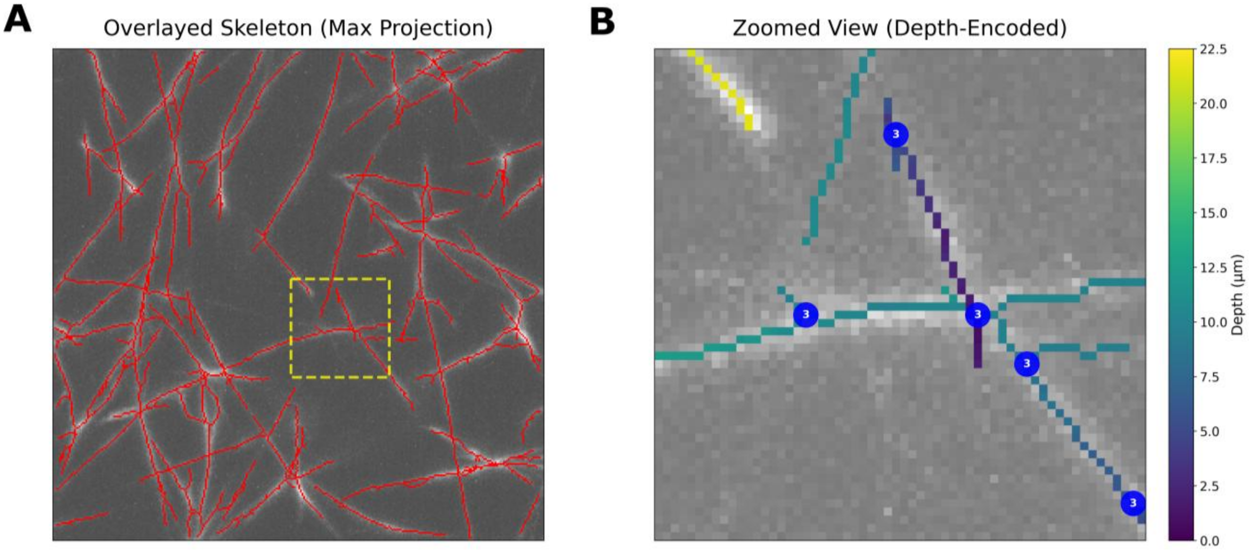


**Supplemental Figure 6: A widened 4-branch junction misidentified as multiple 3-branch junctions.** A) Max z-projection of skeleton with raw volume overlayed and region which will be zoomed into highlighted. B) Zoomed in region with skeleton and junction points marked. Branch number annotated onto each junction point, and depth is displayed by the color of the skeleton segments.

### Simulations:

We built an agent-based model to investigate how fixed-length fibrin fibers diffuse in 3-dimensional space and bind to other fibers to form junctions. The model is initialized with 200 fibers randomly distributed in a cube of volume 115 x 115 x 115 μm with reflecting boundary conditions. The positions of the fiber midpoints are determined by sampling from a uniform random distribution of the simulation domain, minus a boundary margin equal to half the fiber length to ensure that each fiber lays fully inside the cubic domain. Fiber orientations are determined by sampling from a normal distribution in three spatial dimensions. Once the fiber's direction is obtained in this manner, the fiber end points are calculated by extending half the fiber length in the positive and negative orientation from the fiber center.

Fibrin fibers are represented in the model as line segments of length 6 μm. Each line segment is discretized into 20 equally spaced three-dimensional points. Whenever at least one of these 20 points is within a distance of 0.1 μm from a point on another line segment, the fibers connect and form a junction. To accelerate proximity queries, a voxel-based spatial hash is used to identify candidate points lying within a search radius of 0.2 μm from a particular point. Euclidean distances are then computed between the current point and all candidate points. Point pairs whose distance is strictly less than 0.1 μm are potential junctions. If only one pair of points has separation distance less than 0.1 μm, then those are the points that form a connection; if multiple pairs of points satisfy the proximity condition, then a single connection is selected probabilistically as follows. Each candidate pair is assigned a weight proportional to the inverse of its separation distance (dist), $w = \frac{1}{dist + \epsilon}$, where $\epsilon = 1 \times 10^{-10}$ is a small constant to avoid division by 0. The weights are normalized to form a probability distribution, and one candidate is drawn at random according to this distribution, biasing selection toward closer point pairs while still retaining stochasticity. Once a candidate pair is selected, the corresponding points on the two line segments are connected by translating both entire segments so that the two points coincide at their midpoint. This symmetric displacement preserves relative segment geometry while enforcing the connection. The connection is then recorded in a symmetric connectivity matrix to prevent repeated connections between the same segment pair.

Each connection is classified based on whether the selected points lie at segment endpoints or in the interior of the discretization. Endpoint–endpoint connections are labeled as L-junctions, endpoint–interior connections as T-junctions, and interior–interior connections as X-junctions (Supplemental Figure 7). For endpoint–endpoint connections, local tangent vectors at the endpoints are used to compute the interior junction angle from 0-180°. For T- and X-junctions, the overall segment direction vectors (from first to last discretized point) are used. The junction angle is computed as the arccosine of the normalized dot product of the relevant direction vectors. An I-junction is classified as any connection for which the junction angle is either 0 or 180°, however this never occurred in the simulation, leading us to conclude that the experimentally-observed I-junctions must form from a mechanism other than simple diffusion and sticking together.

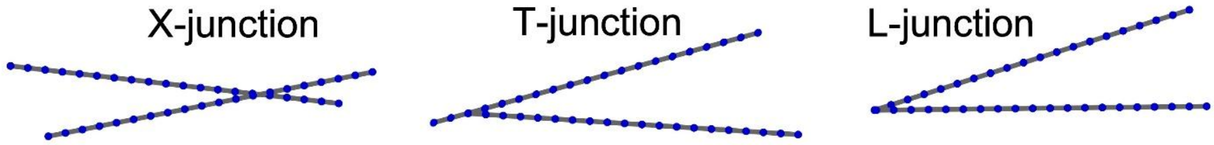


**Supplemental Figure 7: Cartoon examples of different model junction types.** Each line segment represents a fibrin fiber which is discretized into 20 equally-spaced points. An X-junction occurs if two interior discretization points on separate fibers meet; a T-junction occurs if an endpoint of one fiber meets an interior point of a second fiber; an L-junction occurs if the two fibers meet at their endpoints.

We assume that the fibers are rigid, thin rods with diameter 100 nm. Since the diameter is much smaller than the length (6 μm), the translational and rotational diffusion coefficients are calculated using Broersma's corrected slender-body theory, accounting for anisotropic hydrodynamic drag on rigid rods in a viscous solution.(11, 12) The translational diffusion coefficient parallel to the rod axis is given by

$$D_{parallel} = \frac{k_B T}{2\pi\eta L}\left(\ln p - 0.114 - \frac{0.15}{\ln 2p} - \frac{13.5}{(\ln 2p)^2} + \frac{37}{(\ln 2p)^3} - \frac{22}{(\ln 2p)^4}\right).$$

The translational diffusion coefficient perpendicular to the rod axis is given by

$$D_{perp} = \frac{k_B T}{4\pi\eta L}\left(\ln p + 0.866 - \frac{0.15}{\ln 2p} - \frac{8.1}{(\ln 2p)^2} + \frac{18}{(\ln 2p)^3} - \frac{9}{(\ln 2p)^4}\right).$$

The rotational diffusion is given by the equation

$$D_{rot} = \frac{3k_B T}{\pi\eta L^3}\left(\ln p - 0.4466 - \frac{0.2}{\ln 2p} - \frac{16}{(\ln 2p)^2} + \frac{63}{(\ln 2p)^3} - \frac{62}{(\ln 2p)^4}\right).$$

We assume that T = 310.15 K (human body temperature), $\eta$= 1.11474 x $10^{-3}$ kg/(m*s) (viscosity of blood plasma), L = 6.0 x $10^{-6}$ m (length of fiber), $p$ = 60 (aspect ratio), and Boltzmann's constant $k_B$ = 1.380649x$10^{-23}$ ($m^2$*kg)/($s^2$ * K). We multiply the translational diffusion coefficients by $10^{12}$ to convert from $m^2$ to $\mu m^2$ and ultimately obtain model diffusion coefficients of $D_{parallel}$ = 0.3725 $\mu m^2$/s, $D_{perp}$ = 0.2406 $\mu m^2$/s, and $D_{rot}$ = 0.00571 $rad^2$/s.

When fibers connect and form a junction, we assume that the orientation they connected in is permanent; they will remain in this new orientation (relative to each other) for all time, but this new connected component will continue to diffuse and interact with other fibers and components. Because components (2 or more connected fibers) are bigger than a single fiber, they diffuse more slowly. We account for this in the model by slowing down the diffusion coefficients by a factor of 1/(# of fibers in the component)$^{3.}$ For example, a component composed of 4 fibers will diffuse with coefficients that are 1/64 times the diffusion coefficients for single fibers. To account for Brownian motion, each time step we generate random translation and rotation of each fiber and component by sampling from a 3-D Gaussian distribution with variance matching the translational or rotational diffusion, respectively, expected from thermal noise over one time step. A schematic overview of the model is provided in Supplemental Figure 8.

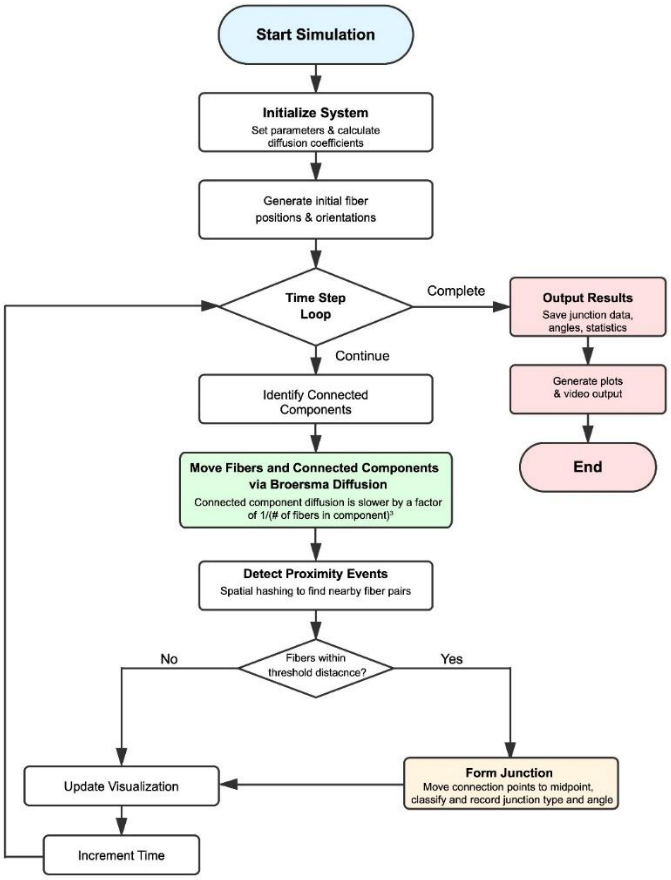


**Supplemental Figure 8: Schematic overview of the computational model for fibrin fiber junction formation.**

### SEM Imaging

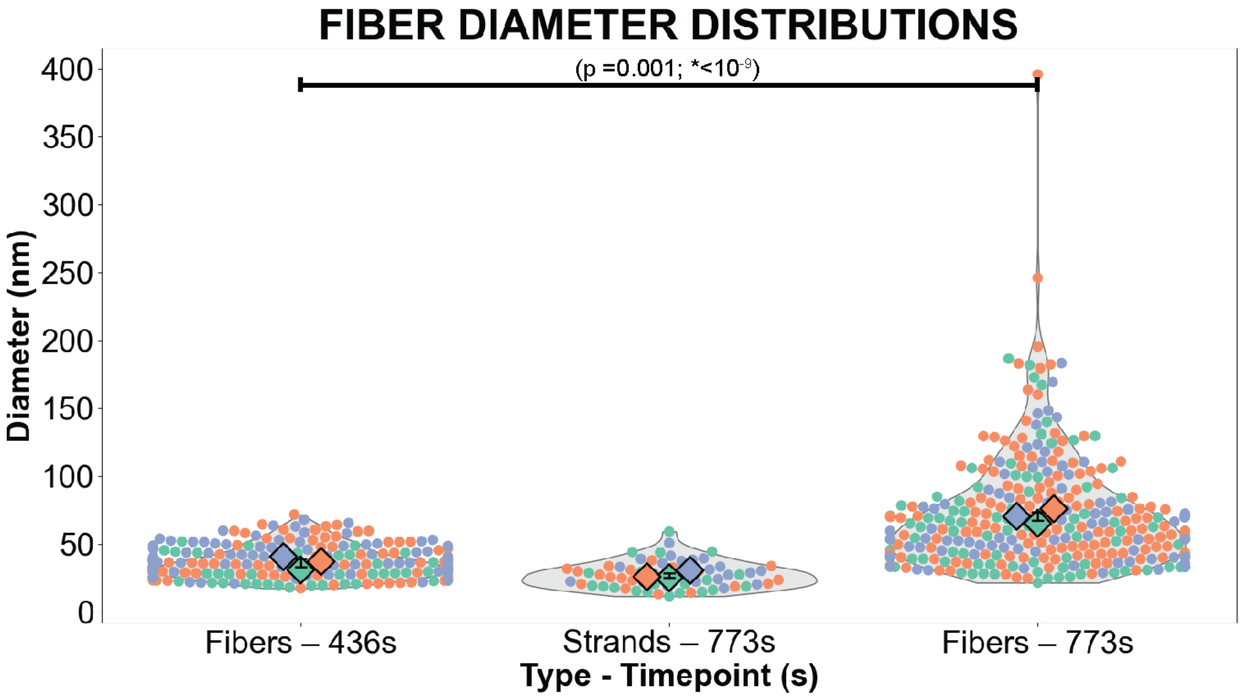


**Supplemental Figure 9. Fiber diameters at early and late timepoints compared with the size of fiber strands.** Data for each timepoint represent N=3 biological replicates, with n=253 (436 s), n=63 (773 s strands), n=313 (773 s fibers). Colored datapoints within each superplot indicate data from a specific biological replicate, with the mean value for each replicate being depicted as a diamond. P-values given for comparison between the mean values (left) and the combined data (*right).

Because SEM coating adds material mainly to exposed surfaces, it inflates the apparent diameter of isolated single strands at 436 s, whereas per-strand measurements within the 773 s cables are less affected (the coating does not substantially change the distance between inter-strand gaps). Thus, the cables at 773 s likely result from lateral bundling of the single-strand fibers present at 436 s.

### Testing the Effect of Fibrinogen Concentration on Fibrin Branching Mechanisms

To determine whether the binding mechanisms changed under different fibrinogen concentrations, we performed additional experiments (N=3) using a final fibrinogen concentration of 3 mg/mL. Under these conditions, similar junction formation event types (X-, T-, L,, and I-) were observed, but the distribution of junction types differed. We performed a Fisher's exact test on the raw counts of X-, T-, L-, and I- junctions observed at 0.5 mg/mL fibrinogen (n = 671) and 3 mg/ml fibrinogen (n = 295). The overall test revealed a significant difference in junction type distributions between the two conditions ($p = 2.3 \times 10^{-6}$). Pairwise Fisher's exact tests with Bonferroni correction identified I- junctions as the primary driver of this difference, with a significantly lower proportion of I- junctions at 3 mg/ml compared to 0.5 mg/mL (9% vs. 23%, p

$= 4.2 \times 10^{-7}$); X- junctions were more prevalent at 3 mg/ml (54% vs. 45%, $p = 0.0096$), though this result should be interpreted cautiously given the Bonferroni-corrected significance threshold of $\alpha = 0.0125$. The proportions of T- and L- junctions did not differ significantly between conditions ($p = 0.114$ and $p = 0.301$, respectively).

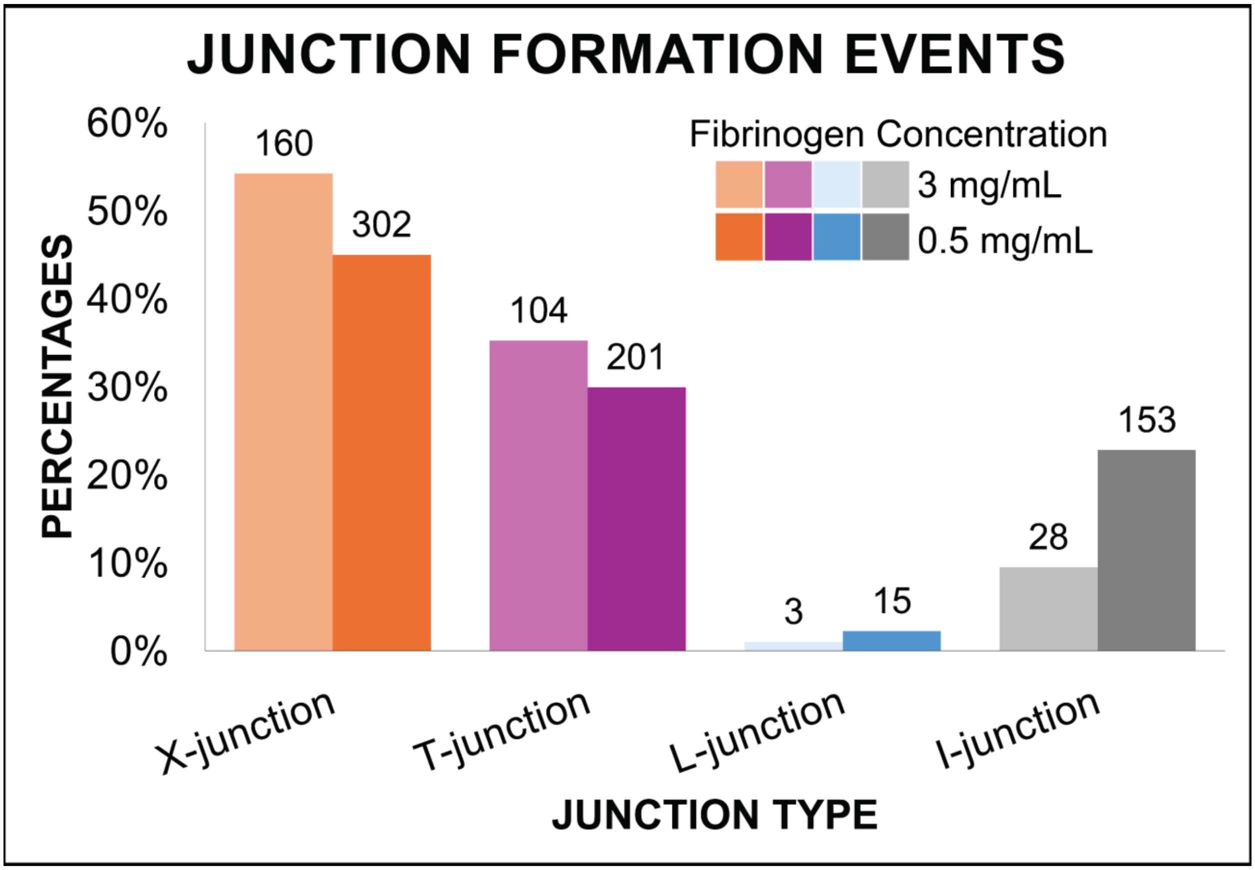


**Supplemental Figure 10. Comparison of fibrin junction formation event percentages at two different fibrinogen concentrations.**

### Supplemental Movie Captions:

Video 1: Maximum intensity projection time series of fibrin polymerization using a modified SiMView light-sheet microscope.

Videos 2-5: Representative movies of X- (Video 2), T- (Video 3), L- (Video 4) and I-junction (Video 5) formation. Blue arrows in initial frame identify fibers that eventually bind. In each movie an initial pan demonstrates nearby but non-bound fibers. Time evolution shows the same fibers diffusing together and binding. A final pan demonstrates that the fibers are bound in 3-D.

Video 6: Simulation of fiber diffusion and junction formation. Line segments represent fibrin fibers and are colored to help distinguish between individual fibers. When two fibers form a junction, a green circle with red outline is plotted to help visualize the location of the junction. All axes are in microns. Each frame of the video corresponds to 10 s of simulation time, for a total of 420 s of simulation time.


1. Tomer R, Khairy K, Amat F & Keller PJ (2012) Quantitative high-speed imaging of entire developing embryos with simultaneous multiview light-sheet microscopy. *Nat Methods* 9(7): 755–763.

2. Lord SJ, Velle KB, Mullins RD & Fritz-Laylin LK (2020) SuperPlots: Communicating reproducibility and variability in cell biology. *Journal of Cell Biology* 219(6): 94.

3. The R Core Team (2021) R: A language and environment for statistical computing. *R Foundation for Statistical Computing*
